# Rules for hardening influenza A virus liquid condensates

**DOI:** 10.1101/2022.08.03.502602

**Authors:** Temitope Akhigbe Etibor, Sílvia Vale-Costa, Sindhuja Sridharan, Daniela Brás, Isabelle Becher, Victor Hugo Mello, Filipe Ferreira, Marta Alenquer, Mikhail M Savitski, Maria João Amorim

## Abstract

Multiple viral infections form biomolecular condensates in the host cell to compartmentalize viral reactions. Accumulating evidence indicates that these viral condensates may be hardened, a strategy with potential for exploitation as novel antiviral therapy, given that viral reactions rely on specific material properties for function. However, there is no molecular understanding on how to specifically and efficiently modify the material properties of viral condensates, a pre-requisite for overcoming off-target effects by rational drug design. *In vitro*, the material properties of biological condensates are modified by different thermodynamic parameters, including free energy, concentration, and type/strength of interactions. Here, we used influenza A virus liquid cytosolic condensates, A.K.A viral inclusions, to provide a proof of concept that modulating the type/strength of transient interactions among the interactome in IAV inclusions is more efficient at hardening these structures than varying the temperature or concentration, both in *in vitro* and in *in vivo* models. This stabilization can be achieved by a known pharmacological sticker that can specifically change the material properties of viral inclusions without affecting host proteome abundance nor solubility. Our work supports the development of antivirals targeting the material properties of biomolecular condensates in viral infections. It also provides a framework for the selection of compounds with this activity for general application and thus provides an advance in disease therapy.

## INTRODUCTION

Central to the spatiotemporal control of reactions in many viral infections is the formation of biomolecular condensates that facilitate key steps of viral lifecycles (Etibor et al., 2021). In influenza A virus (IAV) infection, this is key for assembling its segmented genome, a complex composed of 8 different viral RNA segments (vRNA) (Pons, 1976). Each vRNA is encapsidated by molecules of nucleoprotein (NP) along its length, with one unit of the RNA dependent RNA polymerase (RdRp, consisting of PB2, PB1 and PA) bound to the base-paired RNA termini, forming viral ribonucleoproteins (vRNPs) (Amorim, 2019). How the 8 vRNP complex self-assembles is unknown, although it is known that it relies on RNA-RNA interactions amid vRNPs and is a selective process because most virions contain exactly 8-different segments (Hutchinson et al., 2010). After export from the nucleus where vRNPs are synthetized, vRNPs reach the cytosol and induce the formation of cytosolic condensates, known as viral inclusions (Amorim et al., 2011; Avilov et al., 2012; Chou et al., 2013; Eisfeld et al., 2011; Lakdawala et al., 2014; Momose et al., 2011), which we postulated to be sites dedicated to IAV genome assembly (Alenquer et al., 2019). Interestingly, IAV cytosolic inclusions exhibit liquid properties (fuse and divide, dissolve upon shock and are dynamic) (Alenquer et al., 2019), providing the first indication that defined material properties are critical for the formation of influenza epidemic and pandemic genomes.

As the list of viruses utilizing liquid biomolecular condensates is increasing fast, including reoviruses, human cytomegalovirus, HIV, rabies, measles, SARS-CoV-2 (reviewed in (Etibor et al., 2021; Lopez et al., 2021)), it becomes pertinent to ask whether targeting the material properties could constitute a novel antiviral approach. Recently, the Sonic hedgehog pathway antagonist cyclopamine and its analogue A3E were demonstrated to inhibit human respiratory syncytial virus (hRSV) replication by altering the material properties of viral condensates (Risso-Ballester et al., 2021). However, compounds targeting hRSV-related (Risso-Ballester et al., 2021) and cancer-associated (Klein et al., 2020) condensates exhibited off-target effects. Therefore, a critical advance in condensate disease therapy, including in viral infections, requires the defining of the yet unknown rules for efficiently and specifically targeting selected biological condensates. Such knowledge would create opportunities towards rational design of molecules targeting these structures and hence reduce off target effects. In several studies, it was demonstrated that the properties of biological condensates respond to many factors in a system-dependent manner (Alberti et al., 2019; Falahati and Haji-Akbari, 2019; Hyman et al., 2014; Milovanovic and De Camilli, 2017; Mittag and Parker, 2018; Perdikari et al., 2020; Riback and Brangwynne, 2020; Snead and Gladfelter, 2019). Entropic free energy (Quiroz and Chilkoti, 2015), concentration (Riback et al., 2020), type, number and strength of interactions (Sanders et al., 2020), have been demonstrated to affect the formation and properties of biomolecular condensate. This suggests that *in vivo* strategies able to modify these parameters could offer solutions for drug development. For example, pathways affecting local energy production, consumption or metabolism will alter the free energy landscape of biomolecular condensates (Patel et al., 2017). Similarly, pathways that regulate the local density of condensate drivers could affect concentration (Banani et al., 2016; Riback et al., 2020). Finally, pathways involved in post-translational modifications (Rai et al., 2018), pH (Kroschwald et al., 2018; Munder et al., 2016) or ionic strength (Yang et al., 2020), as well as strategies promoting aggregation or dissolution of condensate interactomes could affect the type, number and strength of interactions (Bracha et al., 2019a; Bracha et al., 2019b; Zhu et al., 2019). However, *in vivo,* it is unknown if changes in free energy, concentration or strength/type of interactions affect equally the material properties and function of biomolecular condensates.

In this work, we meet the critical need to identify the most efficient and specific strategies to harden IAV liquid inclusions. We found that the stabilization of intersegment interactions is more efficient at hardening IAV inclusions than varying the temperature or the concentration of the drivers of IAV inclusions. Importantly, we show that the hardening topological phenotype is observed in the lungs of infected mice. We also report that it is possible to affect viral inclusions without imposing additional changes in host protein abundance and solubility using solubility proteome profiling of infected cells (Sridharan et al., 2022). In sum, our data support the development of strategies targeting the material properties of cellular condensates in viral infections and provides a critical advance in disease therapy.

## RESULTS

### Framework to identify perturbations that harden IAV liquid inclusions

We previously demonstrated that viral inclusions formed by IAV infection display a liquid profile in the sense that they drip, acquire a spherical shape upon fusion and dissolve in response to hypotonic shock or brefeldin A treatment (Alenquer et al., 2019). Here, we identify the best strategies to harden viral inclusions to investigate if altering their material properties may be a novel antiviral therapy. For this, we systematically probed and compared the impact of temperature, concentration, and number/strength of ligations on the material properties of liquid viral inclusions, as a proxy of entropic, molecular and valency contributions, respectively. We selected these parameters as they are well understood to regulate the interactions amongst components and the material properties of condensates (Quiroz and Chilkoti, 2015; Riback et al., 2020; Sanders et al., 2020) (**Figure 1**A). Methodologically, we employed established protocols for the thermodynamic perturbations to directly compare the effect of several parameters in one study. We quantified the impact of these perturbations on the number, nucleation density (ρ=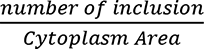, µm^-2^), size, shape, dynamics, supersaturation (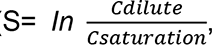, in which C_saturation_ is the concentration above which molecules demix from an homogenous system), and the Gibbs free energy of partition (henceforth called free energy, ΔG = -RT*In*K, in which 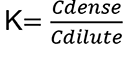 is the partition coefficient). Material concentrations inside (C_dense_) and outside (C_dilute_) viral inclusions were measured using the analytical strategies described in (Riback et al., 2020; Shimobayashi et al., 2021) and shown in **Figure 1**B (and validated in **S3**A-H). For this, we used the mean fluorescence intensity (MFI) of NP as proxy of vRNP concentration (Amorim et al., 2011; Vale-Costa et al., 2016), as it is well established that the majority of cytosolic NP is in the form of vRNPs (Amorim et al., 2011; Avilov et al., 2012; Eisfeld et al., 2011; Momose et al., 2011).

**Figure 1.**
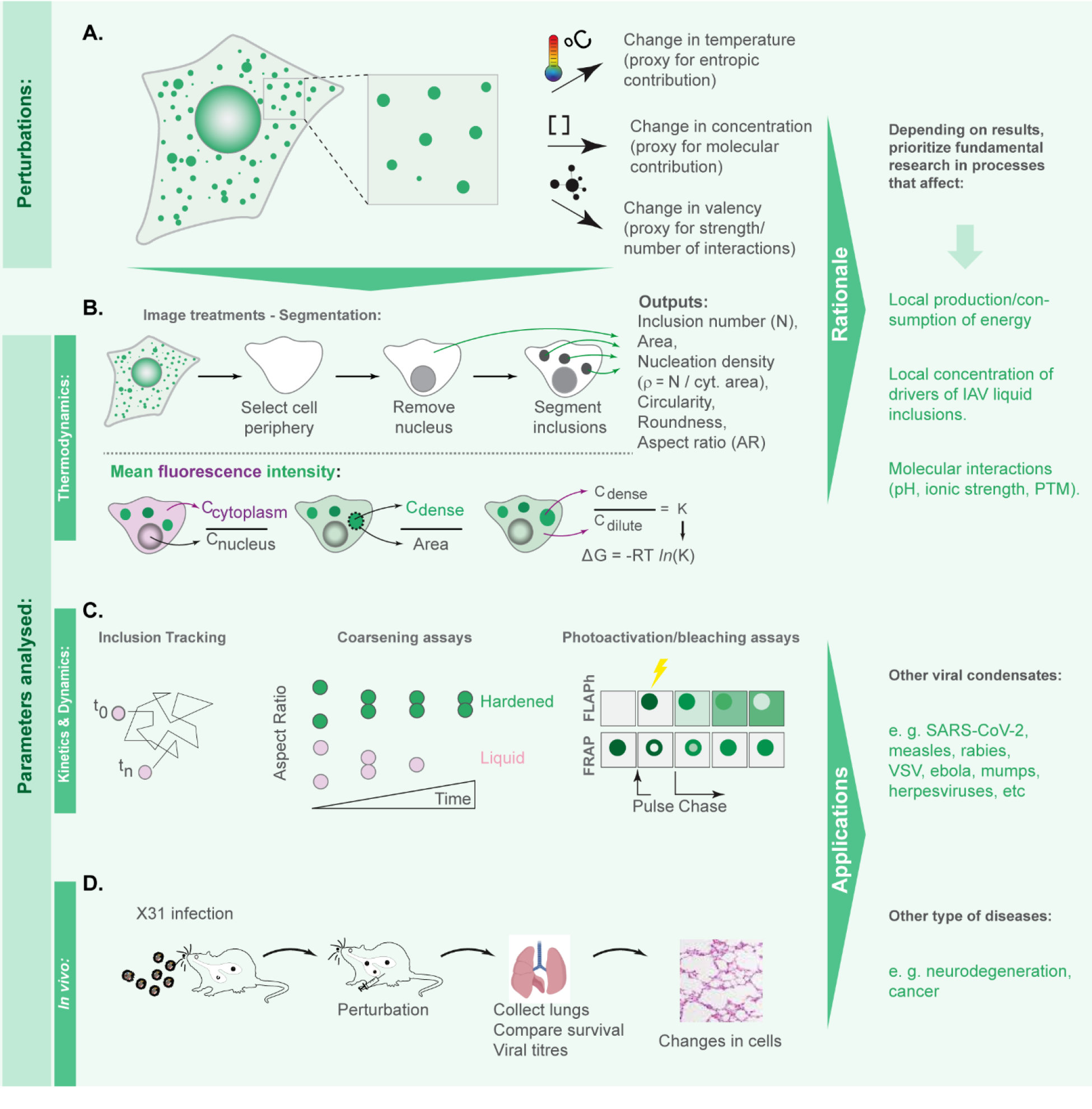
Framework applied to define the rules for hardening IAV liquid inclusions or other condensates. (A) To compare the contributions of entropy, concentration, and valency/strength/type of interactions, we subjected infected cells to the different perturbations, temperature, concentration of viral inclusion drivers (vRNPs and Ras-related in brain 11a (Rab11a)) and number or strength of interactions between different vRNPs using the well-studied vRNP pharmacological sticker, nucleozin. (B) Our aim is to determine which amongst these perturbations impact more dramatically viral inclusions number, shape, size or Gibbs free energy of partition (free energy, ΔG). For this, we segmented circa 20 cells under the different conditions to measure the above-mentioned parameters and the amount of material inside (C_dense_) and outside (C_dilute_) viral condensates. With this, we calculated the partition coefficient K and extrapolated the ΔG. (C) When ΔG dramatically changed, we assessed how perturbations altered the material properties of IAV inclusions by determining how fast and how much they moved (using coarsening assays, particle tracking, fluorescence recovery after photobleaching (FRAP) and fluorescence loss after photoactivation (FLAPh)) (D) We also assessed whether the phenotype could be recapitulated *in vivo* using mice infectied with influenza A virus reassortant X31. The overall goal of this framework is to determine, for IAV, how liquid inclusions may be efficiently hardened to prioritize research and development of strategies with that activity. Additionally, the framework may be applied to other systems, including other viruses, for informed decisions on how to harden condensates.

Our goal was to identify which perturbations translated into significant shifts in ΔG to further explore whether these resulted in dramatic alterations in the material properties of viral inclusions, by assessing their kinetics and dynamics (**Figure 1**C) and determine how they impact viral replication *in vivo* (**Figure 1**D).

### Changes in temperature mildly perturb IAV inclusions

Cellular steady state is maintained at a narrow permissive physiological range, including of temperature. However, biomolecular condensates respond to fluctuations in temperature, and we took advantage of this to assess the entropic contribution of free energy and evaluate whether regulating host cell metabolism could offer future solutions to harden IAV liquid inclusions (**Figure 2**A). For this, we quantitatively analysed the viral inclusions formed in cells incubated at 4 °C, 37 °C and 42 °C for 30 minutes at 8 hours post-infection (hpi) (representative images in **Figure 2**B). Although this short duration in temperature shift is not expected to alter the levels of cytosolic vRNPs, we observed an increase in vRNP amount in the cytosol at 42 °C and a decrease a 4°C (**Figure 2**C). This could be due to the NP antibody having different access to its antigen in IAV inclusions with different morphologies. Increasing the temperature from 37° to 42°C did not significantly change the number (**Figure 2**D-E), size (**Figure 2**G-J) or aspect ratio (**Figure 2**K-L) of viral inclusions but decreased the concentration of vRNPs in condensates (C_dense_) (**Figure 2**F-I, **Table S**1(sheet1)). Importantly, this increase in temperature modestly destabilized the structure, as observed by an increase in Gibbs free energy (-3571 ± 446.1 J.mol^-1^@ 37 °C to -2659.5 ± 398.1 J.mol^-1^@ 42 °C, mean ± SD, **Figure 2 M-O**, **Table S**1 (sheet1)). These alterations are consistent with heat disruption of molecular interactions leading to disassembly of IAV inclusions with temperature. Conversely, decreasing the temperature until 4°C leads to an increase in the number and size of inclusions (shift in area from 0.14 ± 0.027 µm^2^ at 37 °C to 0.2 ± 0.03 at 4°C, **Figure 2D,E**. J and **Table S**1 (sheet1)), as well as in the concentration of vRNPs in inclusions (C_dense_ at 37°C of 44.4 ± 6.66 AU, mean ± sd, and at 4°C of 63.2 ± 6.40 AU, **Table S**1 (sheet1)), and does not significantly change the stability of IAV inclusions as determined by Gibbs free energy (-3658 ± 410.2 J.mol^-1^@ 4 °C, **Figure 2**M-O).

**Figure 2.**
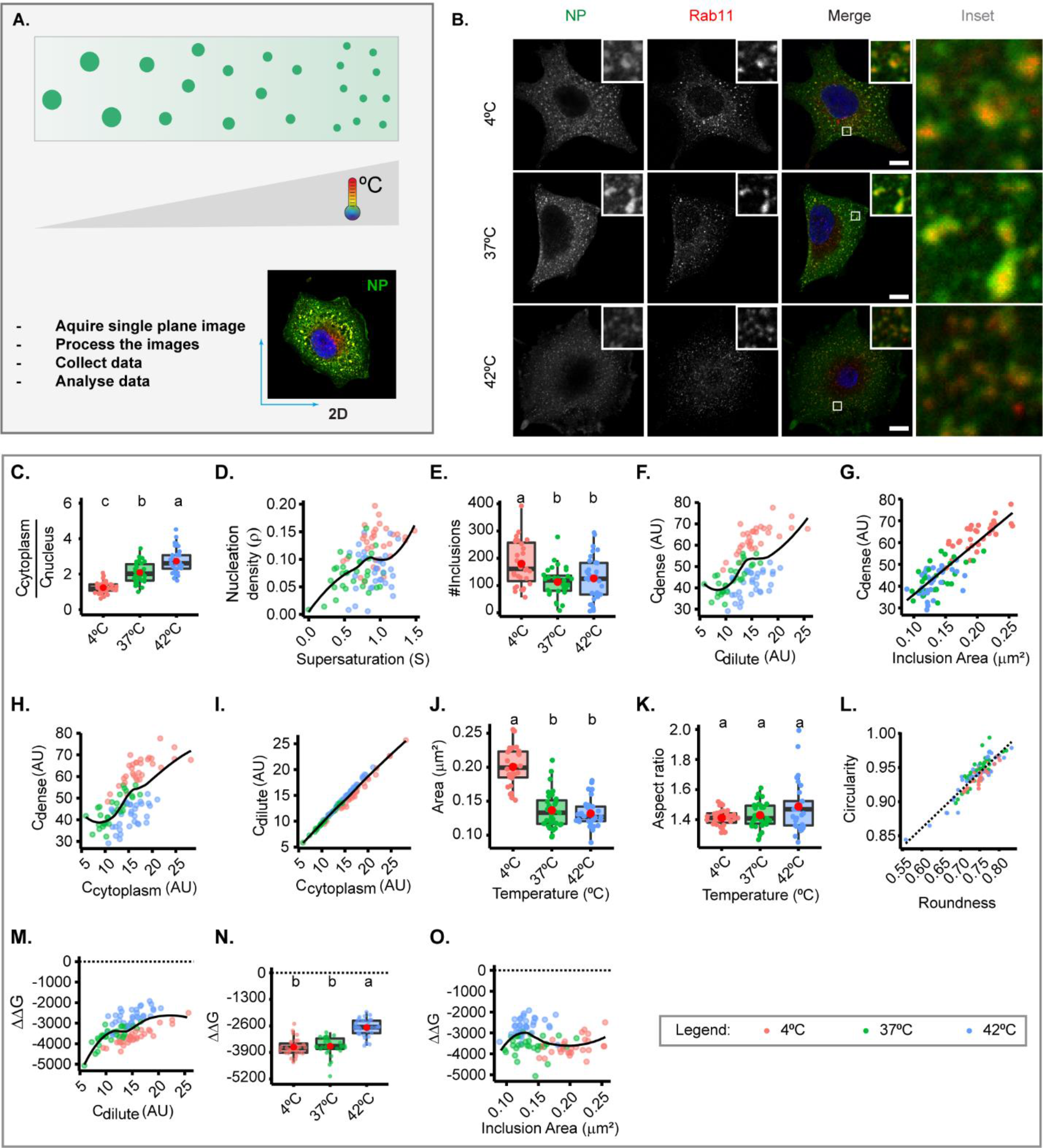
Thermal changes mildly perturb the material properties of inclusions. A549 were infected at a MOI of 3 with PR8 virus for 8 h, incubated at different temperatures (4°C, 37°C, 42°C) for 30 min, fixed, and analysed by immunofluorescence using antibody staining against Rab11 and NP as a proxy for vRNP. The biophysical parameters were extracted from immunofluorescence images (n = 18 - 29), adapting the method published by (Riback et al., 2020; Shimobayashi et al., 2021) to determine concentration Cdense as the mean fluorescence intensity of vRNPs in the segmented IAV inclusions, while concentration Cdilute was extrapolated from the cytoplasmic vRNP intensity outside the inclusions. Each dot is the average value of a measured parameter within or outside IAV inclusions per cell, while the continuous black lines are non-linear fitted models for all data. Also, size and shape of inclusion were extracted from inclusions after image segmentation. Parameters that were normalized to an infection state without IAV inclusions (3hpi) are indicated by a dashed horizontal line. Above each boxplot, same letters indicate no significant difference between them, while different letters indicate a statistical significance at α = 0.05. All data are displayed in **Table S**1 (sheet1). Abbreviations: AU, arbitrary unit. (A) Representative depiction of the experimental analysis workflow. (B) Representative images of fixed A549 cells infected with PR8 virus showing alterations in viral inclusions at different temperatures. (C). Boxplot depicting the fold change in cytoplasmic to nuclear vRNP concentration; *P* = 0.0362 by one-way ANOVA followed by Tukey multiple comparisons of means. (D) Scatter plot of nucleation density 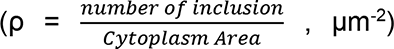 versus degree of supersaturation 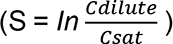, as a measure of propensity to remain dispersed in the cytoplasm. (E) Boxplot showing number of viral inclusions per cell; *P* = 0.00118 by one-way ANOVA, followed by Tukey multiple comparisons of means. (F) Scatter plot of vRNP concentration within inclusions (C_dense_, AU) versus surrounding cytoplasm (C_dilute_, AU). (G) Scatter plot of vRNP concentration in inclusion (C_dense_, AU) versus area of inclusion (µm^2^). (H) Scatter plot of vRNP concentration within inclusions (C_dense_, AU) versus its total cytoplasmic vRNP concentration (C_cytoplasm_, AU). (I) Scatter plot of C_dilute_ (AU) versus total cytoplasmic vRNP concentration C_cytoplasm_ (AU). (J) Boxplot of viral inclusion area (µm^2^) per cell; *P* < 0.00387 by Kruskal Wallis Bonferroni treatment. (K) Boxplot of aspect ratio of inclusions; *P* = 0.234 by Kruskal Wallis Bonferroni treatment. (L) Scatter plot of inclusions circularity versus roundness. (M) Scatter plot of fold change in free energy of partition (ΔΔG, J.mol^-1^) where ΔG = -RT*In*K, and 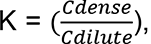, and ΔΔG = ΔG – ΔG_3 hpi_, versus vRNP concentration in the cytoplasm outside viral inclusions (C_dilute_, AU) (N) Boxplot of ΔΔG (J.mol^-1^); *P* < 8.01e-16 by one-way ANOVA followed by Tukey multiple comparisons of means. (O) Scatter plot of relative fold change in ΔΔG versus area of inclusion (µm^2^).

### Changes in concentration of viral inclusions’ drivers do not impact their liquid profile

Two factors were shown to drive the formation of IAV inclusions - vRNPs and Ras-related in brain 11a (Rab11a) (Alenquer et al., 2019; Amorim et al., 2011; Eisfeld et al., 2011; Lakdawala et al., 2014; Vale-Costa et al., 2016; Veler et al., 2022). In fact, vRNP accumulation in liquid viral inclusions requires its association with Rab11a directly via the viral polymerase PB2 (Amorim et al., 2011; Veler et al., 2022), and the liquid character is maintained by an incompletely understood network of intersegment interactions bridging several cognate vRNP-Rab11 units on flexible membranes (Vale-Costa et al., 2016). As the concentration of material is a key determinant for the physical properties of condensates (Hernandez-Vega et al., 2017; Riback et al., 2020; Weber and Brangwynne, 2015), we evaluated how concentration of these two drivers impacts the behaviour of IAV inclusions.

For this, we took advantage of the fact that vRNP levels increase during infection (Kawakami et al., 2011), and we analysed viral inclusions over a time course, in two conditions: with endogenous levels of Rab11a (using cells expressing GFP, **as in (Alenquer et al., 2019)**) and overexpressing Rab11a (in the form of GFP-Rab11a, **as in (Alenquer et al., 2019)**) (**Figure 3**A-B, **Figure S2**). With this approach, we aimed at analysing whether the material properties of viral inclusions changed over time and whether increasing the levels of Rab11 would alter these properties. This strategy would reveal if regulating Rab11a activity could harden IAV liquid inclusions.

**Figure 3.**
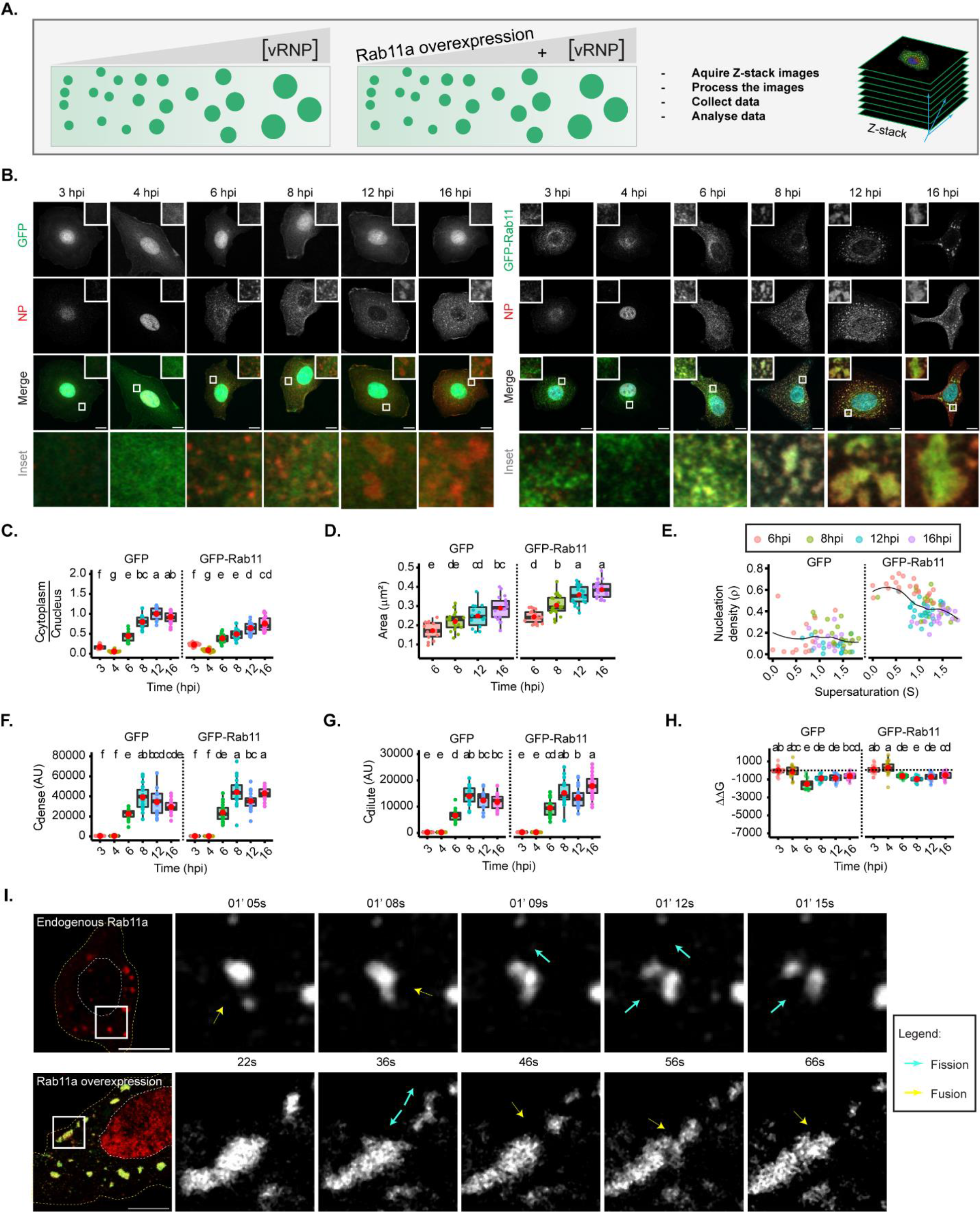
Changes in concentration of vRNPs and Rab11a modestly alter the material properties of viral inclusions. (A - H) A549 cells stably expressing GFP, or Rab11a-WT were infected at a MOI of 3 with PR8 virus and, at the indicated time points, were fixed, and analysed by immunofluorescence using an antibody against NP (as a proxy for vRNPs). (C - H) Each dot is the average value of measured parameters per cell, and the continuous black lines are non-linear fitted models for all data. Above each boxplot, same letters indicate no significant difference between them, while different letters indicate a statistical significance at α = 0.05 using one-way ANOVA, followed by Tukey multiple comparisons of means for parametric analysis, or Kruskal-Wallis Bonferroni treatment for non- parametric analysis. All thermodynamic related values are displayed in **Table S**1 (sheets 2 and 3). Abbreviations: AU, arbitrary unit. (A) Representative depiction of the experimental analysis workflow. (B) Immunofluorescence images of infected cells at different hours post-infection (hpi) (a proxy for changing cytoplasmic vRNP concentration) in cells overexpressing GFP (left) or GFP-Rab11 (right) (both in green); NP (red), and nucleus (blue). Scale bar = 10 µm. (C) Boxplot depicting the fold change in the ratio of cytoplasmic to nuclear vRNPs concentration at different times of infection, with endogenous or overexpressed Rab11a; *P* < 0.001; Kruskal Wallis Bonferroni treatment. (D) Boxplot of inclusion area (µm^2^) per cell; *P* < 0.001 by one-way ANOVA, followed by Tukey multiple comparisons of means. (E) Scatter plot showing nucleation density (ρ, µm^-2^) versus degree of supersaturation (S). (F) Boxplot of C_dense_ (AU); *P* < 0.001 by Kruskal Wallis Bonferroni treatment. (G) Boxplot of C_dilute_ (AU); *P* < 0.001 by Kruskal Wallis Bonferroni treatment. (H) Boxplot of ΔΔG (J.mol^-1^); *P* < 0.001 by Kruskal Wallis Bonferroni treatment. Conditions were normalized to an infection state without IAV inclusions (3 hpi) that is indicated by the dashed black line. (I) A549 cells stably expressing GFP, or Rab11a-WT were transfected with a plasmid encoding mCherry-NP and simultaneously co-infected with PR8 virus at an MOI of 10 and were live imaged at 12 – 16 hpi. Representative time lapse images of fission (blue arrow) and fusion (yellow arrow) dynamics of viral inclusions in cells with endogenous levels or overexpressing Rab11a (**Movie S**1, **S**2).

In GFP expressing cells, as the progeny vRNP pool reaches the cytosol (**Figure 3**A,C), viral inclusions augment in size (from 0.172 ± 0.04 to 0.289 ± 0.06 µm^2^, mean ± SD, **Figure 3**D), with similar aspect ratio (**Figure S1A,B**). There is a mild reduction in the number of inclusions from 8hpi onwards, as measured by the nucleation density (ρ) (**Figure 3**E, **S**1C, all topological data in **Table S**1 (sheet2)). As infection progresses, the concentration of vRNPs inside condensates increases until 8 hpi (**Figure 3**F and **S**1D,E), accompanied by an increase in the diluted cytosolic phase (**Figure 3G** and **S**1D,F, **Table S**1(sheet2)), and both parameters stabilise thereafter, indicating that the critical concentration occurs around 8 hpi. Importantly, Gibbs free energy (normalised to 3 hpi) is lowest at 6 hpi (-1799.0 ± 623 J.mol^-1^) and destabilises mildly onwards (- 1139.8 ± 382, -1131.2 ± 444 and -833.8 ± 342 J.mol^-1^ @ 8, 12 and 16 hpi, respectively) (**Figure 3**H, **S**1G,H, **Table S**1 (sheet2)). These results are consistent with the increase in cytosolic vRNP leading to bigger sized inclusions that overall maintain the same concentration although becoming modestly destabilised, suggesting that the material properties are also modestly affected. When overexpressing Rab11a (right side of each graph), cytosolic vRNPs also accumulated in viral inclusions that increased with infection (**Figure 3**C-D, from 0.243 ± 0.03 to 0.385 ± 0.04 µm^2^), but were significantly bigger than viral inclusions in GFP expressing cells, revealing a higher nucleation density (**Figure 3**E and **S**1C) and similar aspect ratio (**Figure S1A,B**), C_dense_ (**Figure 3**F and **S**1D,E) and C_dilute_ (**Figure 3**G and **S**1D,F). The lowest value of Gibbs free energy occurs at 8 hpi (-1337.7 ± 331 J.mol^-1^) and destabilises from then onwards (-1145.3 ± 443 and -895.3 ± 394 J.mol^-1^ @ 12 and 16 hpi, respectively, **Figure 3**H, **S**1G,H, all thermodynamic data in **Table S**1(sheet3)). This is consistent with Rab11a overexpression giving rise to bigger viral inclusions that overall contained the same vRNP concentration and destabilise slightly later. Importantly, in the two conditions and over the course of infection, viral inclusions maintained a liquid character with fusion and fission events taking place (**Figure 3**I, Movies S1-2). Therefore, these data indicate that altering the concentration of vRNPs and/or Rab11a affects the size but modestly impact IAV inclusions’ material properties.

### The increase in type/strength of vRNP interactions dramatically stabilizes IAV inclusions

Another critical regulator of condensate properties is the type and strength of interactions among ts components interact (Alberti and Hyman, 2021). Therefore, we predict that oligomerizing vRNPs to each other, or to Rab11a, will change the viscoelasticity of condensates in similar manner to iPOLYMER in intracellular hydrogels (Nakamura et al., 2018). For IAV, it was shown by many independent groups that the drug nucleozin operates as a pharmacological sticker that oligomerizes all forms of NP (Amorim et al., 2013; Kao et al., 2010; Nakano et al., 2021). In fact, it was demonstrated that this drug has affinity for 3 different sites in NP (Kao et al., 2010) chemically polymerizing NP either, free or in vRNPs, in a reversible manner (Amorim et al., 2013). Interestingly, nucleozin was described as a novel class of influenza antivirals targeting the viral protein NP, potently inhibiting IAV replication in cultured cells and in a mouse model of influenza infection (Cianci et al., 2012). However, it readily evolved escape mutant viruses carrying the single substitution Y289H in NP (Kao et al., 2010). Despite its capacity to evolve resistance, our strategy is to take advantage of a well-known tool to probe the effects of increasing the number and type of intra and inter-vRNP interactions in the material properties of IAV inclusions (**Figure 4**A).

**Figure 4.**
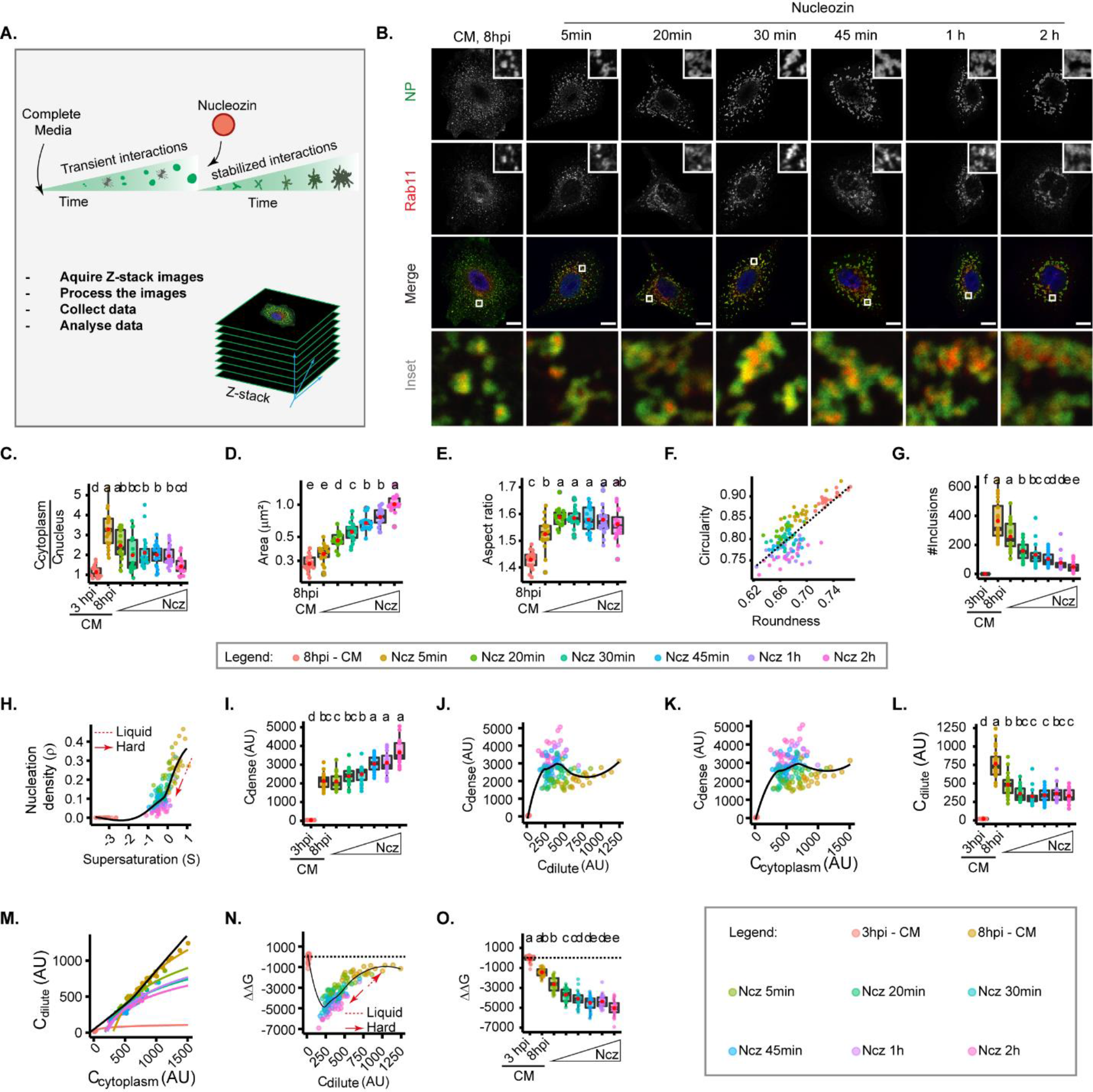
Increasing interaction number and strength stabilizes IAV inclusions. A549 cells were infected at a MOI of 3 with PR8 virus for 8 hrs, then incubated with 5µM of nucleozin (Ncz), a vRNP pharmacological sticker, for different time periods from 5mins to 2 h, before fixing. Cells were processed for immunofluorescence analysis, using antibodies against NP and Rab11a. Each dot is the average value of a measured parameter per cell, while the continuous black lines are non-linear fitted models for all data. Conditions normalized to an infection state without IAV inclusions (3 hpi) are indicated by a dashed black horizontal line. Above each boxplot, same letters indicate no significant difference between them, while different letters indicate a statistical significance at α = 0.05 using one-way ANOVA, followed by Tukey multiple comparisons of means for parametric analysis, or Kruskal-Wallis Bonferroni treatment for non- parametric analysis. All the values calculated for the thermodynamics parameters have been included as **Table S**1 (sheet 4). Abbreviations: AU, arbitrary unit, CM, complete media and Ncz, nucleozin. (A) Representative depiction of the experimental and analysis workflow. (B) Representative images of infected A549 cells subjected (or not) to increasing periods of Ncz treatment. NP (green), Rab11a (red) and nucleus (blue). Scale bar = 10µm. (C) Boxplot depicting the fold change in the ratio of cytoplasmic to nuclear vRNPs concentration before and after Ncz treatment at 8hpi; *P* = 6.16e-14 by Kruskal Wallis Bonferroni treatment. (D) Boxplot of mean inclusion area per cell; *P* < 0.001 by Kruskal Wallis Bonferroni treatment. (E) Boxplot of inclusion aspect ratio; *P* < 2e-16 by Kruskal Wallis Bonferroni treatment. (F) Scatter plot of inclusion circularity versus roundness. (G) Boxplot showing the number of inclusions per cell; *P* < 0.001 by Kruskal Wallis Bonferroni treatment. (H) Scatter plot of nucleation density (ρ, µm^-2^) versus degree of supersaturation (S). (I) Boxplot showing increasing inclusion C_dense_ (AU) with increasing Ncz incubation period; *P* < 0.001 by Kruskal Wallis Bonferroni treatment. (J) Scatter plot of C_dense_ (AU) versus C_dilute_ (AU). (K) Scatter plot of C_dense_ (AU) and C_cytoplasm_ (AU. (L) Boxplot showing C_dilute_ (AU); *P* < 0.001 by Kruskal Wallis Bonferroni treatment. (M) Scatter plot of C_dilute_ (AU) versus C_cytoplasm_ (AU). Coloured lines are non-linear fitted models of grouped data points in the graph. (N) Scatter plot of ΔΔG, J.mol^-1^ versus C_dilute_. (O) Boxplot of fold change in free energy of partition (ΔΔG, cal.mol^-1^); *P* < 0.001; Kruskal Wallis Bonferroni treatment.

With this reasoning, we evaluated the thermal stability of inclusions in the presence or absence of nucleozin in order to confirm its pharmacological sticker activity (Sridharan et al., 2019). It is well established that increasing temperature shifts a thermodynamics system to a homogeneous mix. In agreement, when we exposed IAV infected cells to a range of temperatures (4°C, 37°C and 42°C), we found that higher temperatures yield smaller inclusions tending towards its homogenous distribution in the cytoplasm (**Figure 2**, S2). Interestingly, when infected cells were exposed to the same thermal conditions after nucleozin treatment, inclusions were irresponsive to thermal fluctuation, maintaining their stability (**Figure S2**).

Next, we tracked how nucleozin affected IAV liquid inclusions, by imposing the infected cells to this drug for different periods ranging from from 5 min to 2h. We observes that nucleozin-treated inclusions form a multi-shaped meshwork unlike the rounded liquid droplets formed without nucleozin (**Figure 4**B). Nucleozin affected the concentration of vRNPs in the cytosol that decreased with the time of treatment (**Figure 4**C), presumably by blocking vRNP nuclear export and/or changes accessibility of antibodies to oligomerized NP. Conversely, nucleozin-treatment increased the size of viral inclusions (from 0.284 ± 0.04 without nucleozin to 1.02 ± 0.18 µm^2^ with 2 h treatment, **Figure 4**D), which lost circularity (0.893 ± 0.02 without nucleozin to 0.761 ± 0.02 2h treatment) and roundness (0.734 ± 0.01 without nucleozin to 0.672±0.02 with 2h treatment, **Figure 4**E-F) and decreased in number (from 366.2 ± 133,6 to 48.1 ± 34.0 after 2h treatment **Figure 4**G,H), suggesting that they were stiffer. Interestingly, C_dense_ increased dramatically (from 2125.8 ± 0.09 without nucleozin to 3650.0 ± 0.03 with 2h nucleozin), **Figure 4**I-K) and C_dilute_ decreased and became stable after 20 min treatment (from 766.2 ± 213.0 without nucleozin to 330.2 ± 94.0 after 2h treatment, **Table S**1 (sheet4, total C_dilute_), **Figure 4**J, L-M). Importantly, these structures were energetically more stable, with lower free energy (from -1711.1 ± 397 J.mol^-1^ without nucleozin to -5388.4 ± 808 J.mol^-1^ 2 h post nucleozin addition (**Figure 4**N-O, all topological and thermodynamic values in **Table S**1 (sheet 4)).

Together, the data suggest that stabilizing vRNP interactions changes inclusions more efficiently than the other strategies tested above.

### Modifiers of strength/type of interactions between vRNPs harden liquid IAV inclusions

Changing the strength of interactions amongst vRNPs impacted viral inclusions’ thermodynamics the most. Therefore, we next sought to assess if nucleozin altered their material properties. We first checked if nucleozin-treated viral inclusions maintained the ability to dissolve upon shock treatments, as illustrated in **Figure 5**A. We observed that native inclusions responded to shock treatment as expected, however, nucleozin strongly held inclusions together that did not dissolve when exposed to either hypotonic or 1,6-hexanediol shock treatments (**Figure 5**B, C). This unresponsiveness to shock suggests that IAV inclusions undergo hardening when vRNP interactions are stronger.

**Figure 5.**
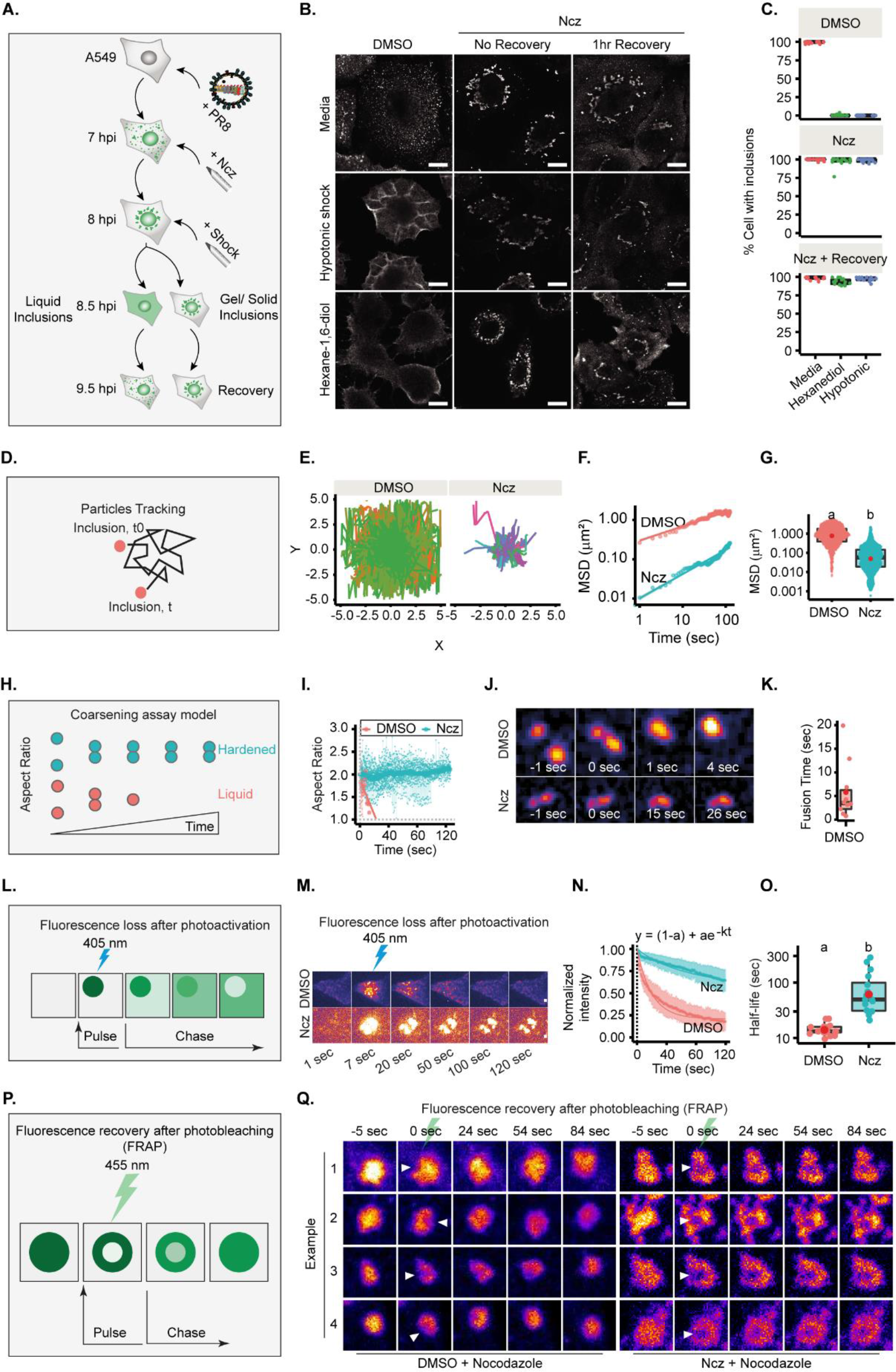
Increasing the strength/type of interactions between vRNPs changes the material properties of liquid IAV inclusions. (A – C) A549 cells were infected at a MOI of 3 with PR8 virus and treated with 5µM Ncz or DMSO at 7hpi. An hour later, cells were treated for 30 min with 80% water (hypotonic shock, Hyp), with 1,6-hexanediol (Hex) or complete media (CM) as control, before allowing recovery from stress treatment in CM for 1 h. Cells were fixed, stained for NP for analysis by immunofluorescence and the percentage of cells with IAV inclusions was scored manually. (D - K, P - Q) A549 cells were infected with PR8 virus at an MOI of 10 and simultaneously transfected with plasmids encoding (D - G) GFP-NP, (H - K) mcherry-NP, or (L -O) photoactivatable GFP-NP and mcherry-NP. Cells were then live imaged after 12 hpi. (A) Experimental schematics of inclusion shock assay. (B) Representative images showing the response of IAV inclusions (NP, as proxy) to shock treatments after incubation in Ncz or DMSO. Scale Bar = 10 µm. (C) Boxplot showing percentage cells with inclusions, after DMSO or Ncz treatment, by manual scoring; P < 0.001 by Kruskal Wallis Bonferroni treatment. (D) Scheme showing how IAV inclusions were tracked over time. (E) Plot showing inclusion (GFP-NP, as proxy) particle trajectory when treated with DMSO or Ncz. (F) Graph showing the mean square displacement (µm^2^) versus time (sec) of IAV. (G) Boxplot depicting the resulting mean square displacement (µm^2^) after 100 sec tracking of IAV inclusions; P < 0.001 by Kruskal Wallis Bonferroni treatment. (H) Schematics of the coarsening assay model, in which liquid and hardened IAV inclusions are represented by orange and blue dots, respectively. Unlike hardened inclusions, native liquid inclusions would fuse and relax to a spherical droplet. (I) Aspect ratio (AR) was used as a measure of IAV inclusion coalescence into a sphere. Mean AR per time was fitted to a linear model (bold coloured lines). Horizontal grey dash lines depict a perfect sphere (aspect ratio = 1). (J) Pseudo-colored time-lapse images of coalescing viral inclusions (GFP-NP used as proxy; extracted from Movie S3,4) in the presence or absence of Ncz. (K) Boxplot of the fusion time (sec) of IAV liquid inclusions. Dots represent fusion time of individual fusion event. (L) Schematic of a fluorescence loss after photoactivation (FLAPh) experiment. (M) Time-lapse pseudo-colour images showing the fluorescence loss in photoactivated IAV inclusions (photoactivatable GFP-NP used as proxy) upon treatment with Ncz or DMSO (extracted from Supplementary Movie S5,6). Bar = 2 µm. (N) Fluorescence intensity decay of photoactivated (PhotoGFP-NP) normalised to the corresponding IAV inclusions expressing mcherry-NP. Coloured lines are single exponential model fitting (y0 = (1-a) + ae^-kt^) of the data point, dots are the mean of the data per second, and vertical lines denote the standard deviation (SD) per time (s). (O) Half-life 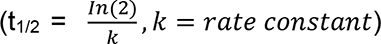 of liquid and hardened IAV inclusions decay post-activation (sec); P = 1.386e-6 by Kruskal Wallis Bonferroni treatment. (P) Schematic depiction of an internal rearrangement of viral inclusion after a ROI within the inclusion is FRAPed. (Q) A549 cells were transfected with plasmids encoding mcherry-NP and co-infected with PR8 virus at an MOI of 10. At 12hpi, cells were treated with nocodazole (10 µg/mL) for 2h to reduce the highly stochastic motion of liquid IAV inclusions and subsequently treated with DMSO or Ncz. Small regions inside IAV inclusions were photobleached to assess internal rearrangement of vRNPs (mCherry-NP as proxy). Time lapse pseudocolor images shows the fluorescence recovery after photobleaching (FRAP, extracted from Supplementary Videos 7,8).

To formally establish that IAV liquid inclusions can be hardened, we compared the dynamics of viral inclusions in the presence or absence of nucleozin using four different approaches. First, we assessed their movement and measured speed and displacement from their point of origin (**Figure 5**D). Native liquid inclusions (treated with sham vehicle - DMSO) display a highly stochastic movement and long displacement, whilst nucleozin-hardened inclusions were less mobile with smaller displacement, as observed by analysing loss of movement in individual tracks (**Figure 5**E). There is an overall reduction in mean square displacement (MSD) with nucleozin (**Figure 5**F) that results in a lower MSD at 100 sec (MSD_100sec_ = 0.838 ± 1.17 µm^2^ without nucleozin shifting to 0.057 ± 0.22 µm^2^ with treatment, median ± SD, **Figure 5**F-G and **Table S**1 (sheet 5)). In a second approach, we measured the time that two droplets take to relax to a sphere upon fusion by coarsening assays (shifting the aspect ratio from 2 to 1, **Figure 5**H). DMSO-treated inclusions relax fast to a single sphere upon fusion (5.8 ± 1.94 s; mean fusion time ± SEM), shifting the aspect ratio from 2 to 1. Nucleozin-treated inclusions retain a stable aspect ratio over time (**Figure 5**I), as they are unable to fuse (**Figure 5**I-K, **Table S**1 (sheet 6), **Movie S**3,4). The results demonstrate that nucleozin stiffens IAV inclusions.

In a third approach, inclusion molecular dynamics was tested by Fluorescence Loss After Photoactivation (FLAPh, **Figure 5**L). In a live imaging experiment, a region of interest (ROI) was photoactivated (**Figure 5**M), its decay profile monitored for 120 sec and the plot fitted to a single exponential model. DMSO- and nucleozin-treated inclusions exhibited distinct decay profiles (**Figure 5**N), with half-life of 14.41 ± 0.9 s (mean ± SEM) and 85.02 ± 19.8 s, respectively (**Figure 5**O, **Table S**1 (sheet 7) and **Movie S**5,6). This indicates that nucleozin treated inclusions become more static.

Lastly, we measured the internal rearrangement in viral inclusions. Because of the small size and highly dynamic nature of IAV inclusions, previous attempts to perform Fluorescence Recovery After Photobleaching (FRAP) experiments resulted in highly variable recovery rates (Alenquer et al., 2019; Amorim et al., 2011) that were unable to accurately determine if internal rearrangements were taking place viral inclusions. As the microtubule depolymerising drug nocodazole largely blocks the movement of IAV inclusions, rendering them larger and more spherical (Amorim et al., 2011; Avilov et al., 2012), we opted for bleaching IAV inclusions upon treating them with nocodazole (**Figure 5**P). In native conditions, the photobleached region quickly disappeared, consistent with internal rearrangement of vRNPs inside IAV inclusions, whilst in nucleozin-treated inclusions, the photobleached area remained unaltered, revealing stiffness (several examples in **Figure 5**Q and **Movie S**7,8).

Taken together, DMSO- and nucleozin-treated IAV inclusions exhibit distinct responses to shocks, dynamics, internal rearrangement and coalescing properties, supporting that nucleozin hardens IAV liquid inclusions.

### Modifiers of strength/type of interactions between vRNPs hardens IAV liquid inclusions *in vivo*

Recently, the condensate-hardening drugs steroidal alkaloid cyclopamine and its chemical analogue A3 were shown to reduce viral titres in respiratory syncytial virus (RSV) infected mice (Risso-Ballester et al., 2021). However, at the organismal level, it was not demonstrated that RSV inclusion bodies in infected cells retained hardened features. To test if we could phenocopy the *in vitro* function of nucleozin, we aimed at analysing vRNP morphology inside the lung cells of infected mice. For this, we challenged mice with the IAV strain X31 for 2 days. At 30 min, 1 h or 2 h before the collection of the lungs, each mouse was treated with PBS (sham vehicle) or nucleozin, administered intranasally (**Figure 6**A-C). Interestingly, when we analysed viral inclusions under control conditions in cells of lungs of infected mice, we observed a punctate-like NP distribution. Upon nucleozin treatment, these cytosolic inclusions grew larger (inclusions per cell mean ± SEM: Ncz 30 min, 0.101 ± 0.006 µm^2^; 2h, 0.226 ± 0.012 µm^2^, **Figure 6**B, **Table S**1 (sheet 8)). This indicates that the pharmacological induced sticker activity of nucleozin (Amorim et al., 2013; Kao et al., 2010) was retained *in vivo.* Having seen an effect in vRNP cytosolic localization in vivo, we aimed at confirming a nucleozin-dependent abrogation of IAV infection in our system as reported before (Kao et al., 2010). In fact, nucleozin was reported to affect viral titres by 1 log and increase survival of IAV (A/Vietnam/1194/04 H5N1) infected mice by 50%. For this, we therefore challenged nucleozin pretreated mice with X31 and treated with a daily dose of PBS (sham vehicle) or nucleozin and found that nucleozin-treated mice had a faster recovery from viral infection (**Figure 6**D). In sum, the data serves as proof of concept that the material properties of condensates may be targeted *in vivo*.

**Figure 6.**
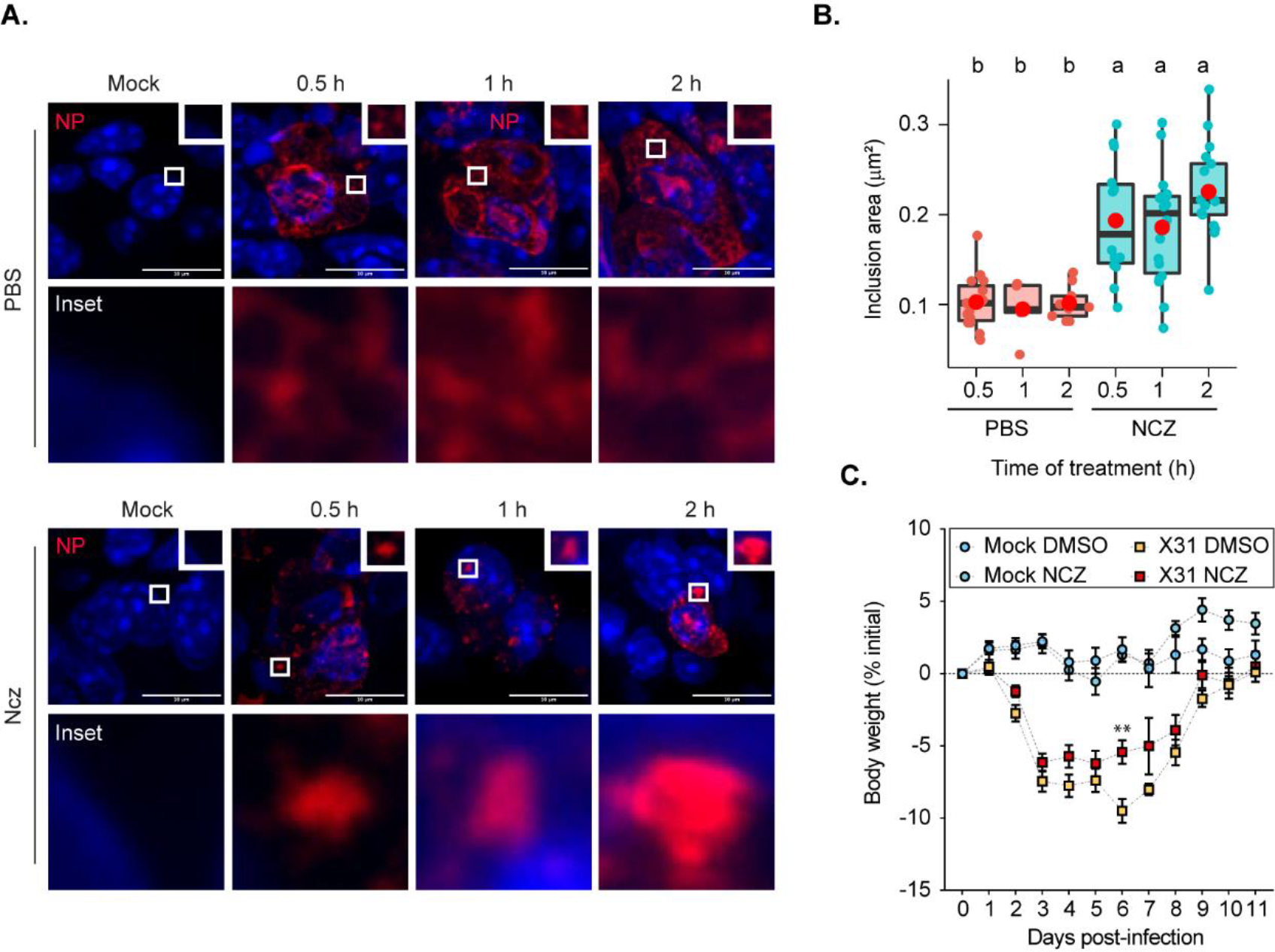
Hardened inclusions emerge *in vivo* when infected mice are treated with nucleozin. (A - B) Mice were intranasally infected with 4000 plaque forming units (PFU) of X31 virus, and after 2 days were intraperitoneally injected with PBS or 8.3 nmoles/g mice of Ncz at 30 min, 1h or 2h before the collection of the lungs. Data were extracted from inclusions (NP, as proxy) from fixed immunofluorescence images of lung tissues. (A) Representative immunofluorescence images show sections of lung tissue stained for NP (red) and nucleus (blue) after PBS or Ncz treatment. (B) Boxplot showing the mean area (µm^2^) of inclusions from cells in lung section;. *P* = 3.378e-8 by Kruskal Wallis Bonferroni treatment. (C) Mice were pre-treated intraperitoneally with 8.3 nmoles/g mice Ncz or PBS for 1 h before being intranasally infected with 1000 plaque forming units (PFU) of X31 virus, injected with a daily dose of Ncz or PBS for 11 days and the weight loss monitored daily.

### Nucleozin rescues formation of hardened IAV inclusions in the absence of Rab11a

Given the possibility to harden IAV inclusions, it is important to define the molecular mechanisms conferring the material properties of these condensates, which remain elusive. As Rab11a drives the formation of IAV inclusions (Alenquer et al., 2019; Amorim et al., 2011; Eisfeld et al., 2011; Lakdawala et al., 2014; Vale-Costa et al., 2016; Veler et al., 2022), we asked if nucleozin could artificially reform viral inclusions and mimic its behaviour in the absence of Rab11a. Stable cell lines expressing Rab11a dominant negative (DN) (henceforward Rab11a-DN) did not form IAV inclusions, as expected, maintaining vRNPs dispersed throughout the cytosol (**Figure 7**A). Interestingly, both Rab11a-WT and Rab11a-DN cell lines, in the presence of nucleozin, exhibited cytosolic puncta (despite smaller in Rab11a-DN lines, (**Figure 7**A-B)). This indicates that nucleozin bypasses the need for Rab11a to concentrate vRNPs, forming aberrant inclusions as predicted. We next tested the fusion ability of nucleozin-induced IAV inclusions in Rab11a-DN cell lines. Unlike native inclusions in WT cells, nucleozin-induced IAV inclusions in Rab11a-DN infected cells are not able to fuse in coarsening assays (**Figure 7**C-E). In sum, the liquid properties of IAV inclusions derived from flexible intersegment interactions and interaction with Rab11a harden to form stiff aggregates upon nucleozin treatment even when active Rab11a is absent.

**Figure 7.**
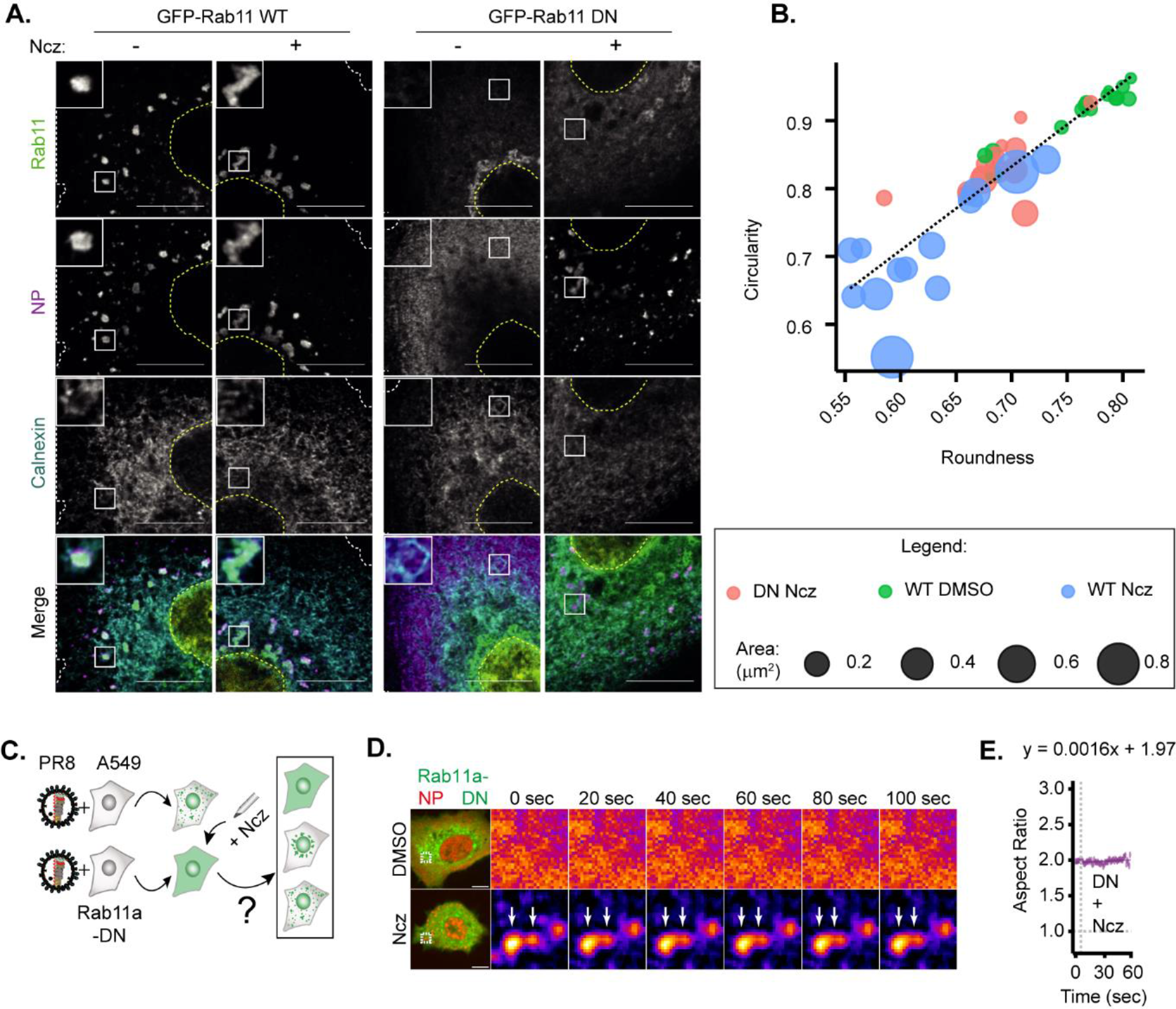
Only hardened inclusions emerge in nucleozin-treated Rab11a-DN cell line. (A - B) A549 cells constitutively expressing GFP-Rab11a-WT and GFP-Rab11a-DN were infected for 10 h with PR8 at a MOI of 3 and treated with 5µM Ncz or DMSO before fixing for analysis by immunofluorescence. (A) Representative images of cells analysed by immunofluorescence staining using antibodies against viral protein NP (magenta), host Rab11 (green) and ER (cyan). Nuclei and cell periphery delimited by yellow and white dashed line respectively, and white boxes are insets showing presence or absence of viral inclusions. Scale bar = 10 µm. (B) Scatter plot of circularity versus roundness of viral inclusions. (C - E) A549 cells constitutively expressing GFP-Rab11a-DN were transfected with mcherry-NP and co-infected with PR8 virus at a MOI of 3. At 12hpi, the cells were treated with 5µM Ncz or DMSO for 10 mins before imaging. (C) Schematic depicting the possible outcomes when Rab11a-DN cell lines are treated with Ncz. (D) Time lapse pseudocolor images show fusion of IAV inclusions in a coarsening assay of PR8 infected Rab11a-DN cell line treated with Ncz or DMSO (extracted from Supplementary Videos 9,10). (E) Plot depicting the aspect ratio of fusing inclusions over time in infected Rab11a-DN cell line treated with Ncz.

### Nucleozin affects vRNP solubility in a Rab11a-dependent manner without altering host proteome profile

Next, to understand how both the viral and host proteomes remodel in response to nucleozin treatment, we used a recently developed quantitative mass spectrometry-based approach called solubility proteome profiling (SPP) (Sridharan et al., 2019). This is a lysate centrifugation assay, which can distinguish the soluble (supernatant) from insoluble (dense assemblies) protein pools. The majority of proteins annotated to be part of membraneless organelles, as well as many cytoskeletal proteins, exhibit prominent insolubility. In SPP, two aliquots of cellular lysates are extracted with either a strong (SDS) or a mild (NP40) detergent. Protein extracted with SDS represent the total proteome, while the supernatant of NP40-extracted lysate represents the soluble sub-pool. The ratio of NP40- and SDS-derived protein abundance represents the solubility of a protein (**Figure 8**A). Protein solubility is a proxy to track phase transition events in different cellular states. However, this measurement cannot distinguish between different events, such as solidification, phase separation, percolation and gelation (Alberti and Hyman, 2021) that may underlie the phase transition.

**Figure 8.**
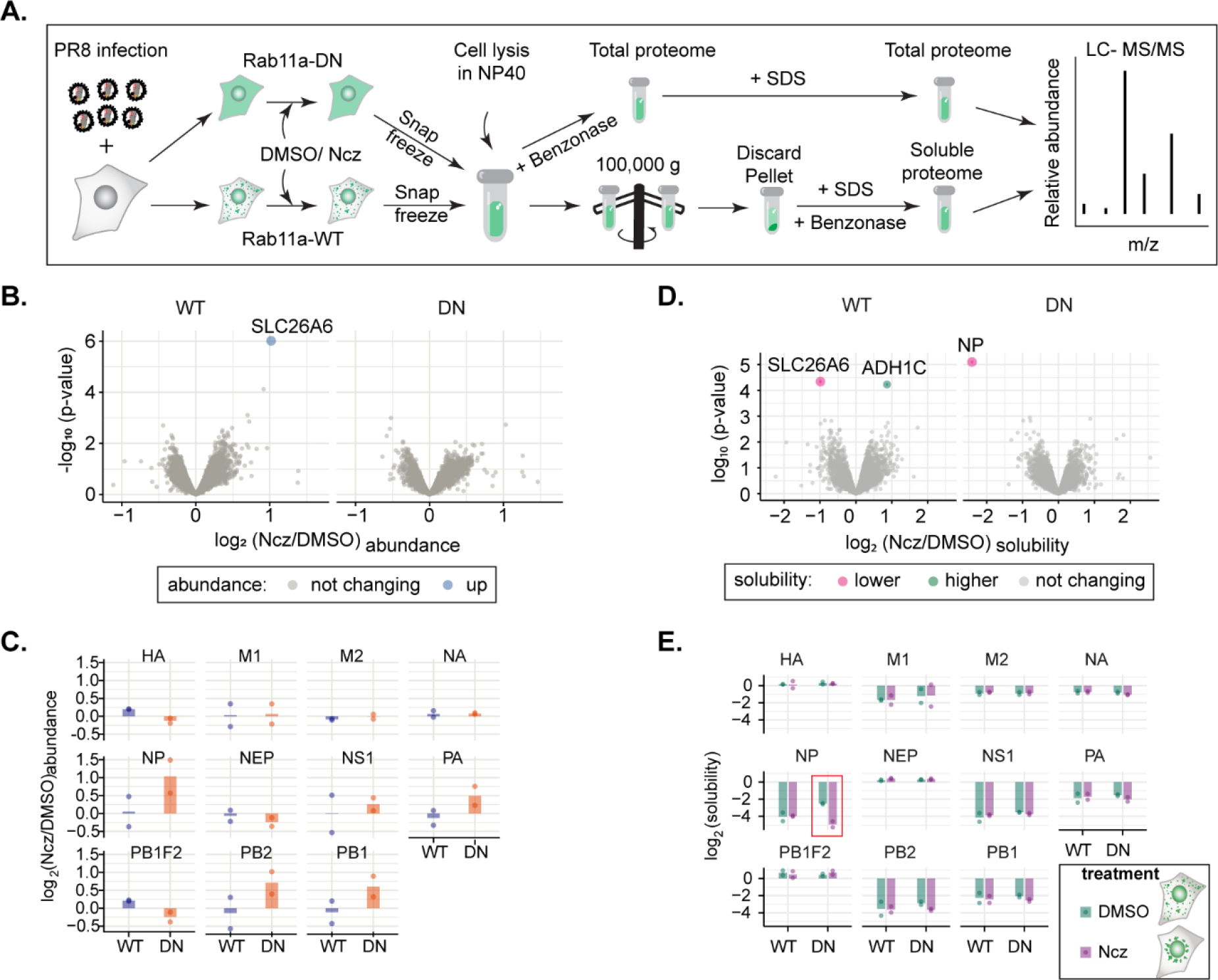
Hardening of IAV inclusions changes its proteome solubility. (A - E) A549 cells constitutively expressing GFP-Rab11a-WT or GFP-Rab11a-DN were infected for 12 h with PR8 at a MOI of 5 and treated with 5µM Ncz or DMSO for 1 h. Thereafter, cells were lysed in mild (NP40) or strong detergent (SDS), while NP40 lysate was ultracentrifuged (100,000 g) to pellet materials in condensates from the soluble fraction in the supernatant. Soluble and total host and viral proteome were identified by LC-MS/MS and solubility was determined as the ratio of soluble NP40- to SDS- derived total proteome abundances at the indicated time points. (A) Schematic representation of solubility proteome profiling (SPP). (B) Volcano plot representing relative host protein abundance in Rab11a-WT and Rab11a-DN infected cell lines (at 12 hpi) after treatment with Ncz or DMSO. Differentially upregulated proteins in these conditions (statistical significance – *see methods*) are indicated in blue dots (C) Bar graphic comparing abundances of viral proteins (in log_2_ scale) in Rab11a-WT and Rab11a-DN cell lines PR8-infected (12 hpi) and treated with either Ncz or DMSO. (D) Volcano plot representing relative solubility of host and viral proteins in Rab11a-WT and Rab11a-DN infected cell lines (at 12 hpi) after treatment with Ncz. Differentially soluble proteins in these conditions (statistical significance – *see methods*) are indicated in pink and green dots. (E) Bar graph comparing solubility (in log_2_ scale) of viral proteins when PR8 infected (12 hpi) Rab11a-WT and Rab11a-DN cell lines were treated with either Ncz or DMSO.

To define the effect of nucleozin in viral inclusions, we compared proteome abundance and solubility profiles of Rab11a-DN cell lines, where the formation of liquid inclusion is blocked, with that of Rab11a-WT cell lines at 12 hpi, in the absence or presence of nucleozin (1 h treatment) (**Figure 8**A-E, **Table S**2,3). Nucleozin-treatment did not induce significant alteration in host proteome abundance in both cell lines (**Figure 8**B). Similarly, no major changes in terms of protein solubility were observed for the host proteome during this treatment period (**Figure 8**B). Overall, our results suggest that nucleozin does not induce changes in cellular protein levels or their solubility.

In terms of the viral proteome, the abundance of all protein components of vRNPs (NP, PB1, PB2, PA and M1) show a modest increase in Rab11a-DN cell lines (**Figure 8**C). On the solubility level, NP exhibited a prominent change. NP remains more soluble in Rab11a-DN lines compared to Rab11a-WT infected cells (fold change of 0.188, *P* = 7.97E-6, **Figure 8**C). This corroborates the observation that vRNPs remain uniformly distributed in Rab11a-DN cells. Upon nucleozin treatment, SPP data reveal that the solubility of NP remains unaltered in Rab11a-WT cells, while increasing the proportion of NP in insoluble pool in Rab11a-DN cells (**Figure 8**D-E, red square). Although there were no changes in solubility by SPP, we observed IAV inclusions growing larger and hardening upon nucleozin treatment at the microscopic level in Rab11a-WT cells (**Figure 7**A). This can be explained, as vRNPs are already insoluble in viral inclusions before nucleozin treatment and the net increase in size of the inclusions does not result in higher insolubility of vRNPs. Both SPP and microscopy complement each other in the case of Rab11a-DN cells, as viral inclusions change from soluble to insoluble and become bigger upon nucleozin treatment. Overall, these data substantiate our finding that vRNPs form Rab11a-dependent insoluble and liquid inclusions that undergo a distinctive (aberrant) phase transition upon nucleozin treatment.

## DISCUSSION

In thermodynamics, the demixing from the surrounding media implies a preference of alike molecules to interact and self-sort, excluding the milieu. This is well understood for binary systems but deviate considerably for muticomponent systems, even *in vitro* (Klosin et al., 2020; Riback et al., 2020; Snead et al., 2022). How living cells, that are complex multicomponent systems at non-equilibrium, operate lacks understanding. Small alterations in the interactions, caused by changes in the environment or the interactome of the condensate, originate different self-assembled structures (Riback et al., 2020) that respond distinctly to thermodynamic variables such as concentration, temperature and type/strength of interactions. For example, increasing the concentration in a system is mostly associated with more ordered, less flexible structures, however higher ordered structures were reported to arise in response to a concentration reduction (Helmich et al., 2010). Therefore, understanding how physical modulators of phase transitions impact the properties of condensates is key to comprehend how biological systems may be regulated, which is essential for, for example, designing condensate-targeting drugs with specific activities (Hermans et al., 2009). IAV infection forms cytosolic liquid inclusions that are sites for genome assembly. Our study to address the fundamental question of whether the material properties of IAV inclusions may be modulated, shows that IAV inclusions may be hardened by targeting vRNP interactions but not by lowering the temperature down to 4 °C nor by altering the concentration of the factors that drive their formation. The data on temperature reveals that a decrease in the entropic contribution leads to a growth of condensates, as observed for other systems (Falahati and Haji-Akbari, 2019; Hyman et al., 2014; Riback et al., 2020), that is, however, mild and does not significantly impact the stability of the structures. Similarly, altering the concentration of drivers of IAV inclusions impact their size but not their material properties. This is unexpected because many studies have shown that changing the temperature or concentration of condensate drivers dramatically impacts their phase diagrams (Bracha et al., 2018; Riback et al., 2020; Zhu et al., 2019) and material properties (Shin et al., 2017). For influenza, these minor effects demonstrate that is system is flexible, which may result from the necessity to maintain the liquid character over a wide range of vRNP concentration in the cytosol (low levels in the beginning and high at late stages of infection). The maintenance of the liquid character may be a regulated process involving fission and fusion events associated with the ER, as reported for other systems (Lee et al., 2020). In fact, IAV liquid inclusions develop in proximity to a particular part of a modified endoplasmic reticulum (ER) (de Castro Martin et al., 2017), the ER exit sites (Alenquer et al., 2019). In addition, the fusion and fission events of inclusions may be necessary to promote vRNP interactions, which is essential for genome assembly, as proposed before (Eisfeld et al., 2015; Lakdawala et al., 2014).

Defining the rules for hardening the condensates is important for understanding how biological condensates may be manipulated in cells and has consequences for development of novel antiviral treatments. By demonstrating that targeting the type/strength of interactions modulates the material properties of liquid viral inclusions in *in vitro* and *in vivo* models, we show that the development of molecules that affect the interactions between two components (such as post-translational modifications, local pH or ionic strength or pharmaceutical stickers/spacers) should be prioritized over those increasing their concentration or local entropy. Such targeting may prevent off-target effects, especially by developing compounds able to distinguish free vRNP components from those in the supramolecular complex. In fact, the solubility proteome profiling herein reported demonstrates that it is possible to harden a liquid condensate without imposing changes in the host proteome abundance and solubility, which is important to increase specificity. However, a cost of targeting conserved molecules is the evolution of escape mutants (Cheng et al., 2012; Hu et al., 2017; Kao et al., 2010). Therefore, a concern to address in the future is how to design suitable combinatory therapies able to reduce their emergence. Since single nucleotide mutations underpin numerous resistance mechanisms to antivirals (Lampejo, 2020), an alternative is to engineer condensate hardening drugs that require multiple amino acid changes for escaping.

In this work, we explored the rules for hardening IAV liquid condensates. Other alternatives to modulate the material properties tailored for function can be developed. For example, accumulating evidence shows that blocking viral inclusion formation hinders viral infection (Amorim, 2019; Amorim et al., 2011; de Castro Martin et al., 2017; Eisfeld et al., 2011; Han et al., 2021; Momose et al., 2011; Vale-Costa et al., 2016; Vale-Costa and Amorim, 2017; Veler et al., 2022). Herein, we observe that increase in temperature biases the system to dissolving viral inclusions, therefore activating exothermic reactions close to IAV inclusions may lead to their dissolution. Furthermore, it has been previously demonstrated that blocking Rab11 pathway, directly or indirectly, hampers viral infection (Amorim et al., 2011; Eisfeld et al., 2011; Han et al., 2021; Momose et al., 2011). Future research could also explore this route. As Rab11a has emerged as a key factor for the replication of members of many unrelated viral families relevant for human health (*Bunyaviridae*, *Filoviridae*, *Orthomyxoviridae*, *Paramyxoviridae* and *Pneumoviridae*), targeting its activity may serve as a pan-antiviral strategy (Amorim et al., 2011; Bruce et al., 2010; Cosentino et al., 2022; Nakatsu et al., 2013; Nanbo and Ohba, 2018).

## Limitations of the study

Understanding condensate biology in living cells is physiological relevant but complex because the systems are heterotypic and away from equilibria. This is especially challenging for influenza A liquid inclusions that are formed by 8 different vRNP complexes, which although sharing the same structure, vary in length, valency, and RNA sequence. In addition, liquid inclusions result from an incompletely understood interactome where vRNPs engage in multiple and distinct intersegment interactions bridging cognate vRNP-Rab11 units on flexible membranes (Chou et al., 2013; Gavazzi et al., 2013; Haralampiev et al., 2020; Le Sage et al., 2020; Shafiuddin and Boon, 2019; Sugita et al., 2013). At present, we lack an *in vitro* reconstitution system to understand the underlying mechanism governing demixing of vRNP-Rab11a-host membranes from the cytosol. This *in vitro* system would be useful to explore how the different segments independently modulate the material properties of inclusions, explore if condensates are sites of IAV genome assembly, determine thermodynamic values and thresholds accurately and validate our findings. One of the constraints of using cells in this work relates to the range and precision of the concentrations we can vary in our system. Herein, we compared endogenous Rab11a cellular levels to a single pool of transduced cells that contained low, but still heterogeneous, levels of Rab11a as a way to avoid toxicity and/or uncharacterized effects of overly expressing Rab11a in the cell. To minimize this limitation, we combined overexpressing Rab11a with a range of low and high levels of vRNPs (analysing the entire time course of infection) to understand if a combination of high levels of vRNPs and of Rab11a could synergistically change the material properties of IAV inclusions. Finally, technically we retrieved thermodynamic parameters (such as C_dense_, C_dilute_, shape, size) from images in z-stacks as the sum of slices at specific snapshots of infection. However, although requiring a very complex imaging analysis that we lack, in the ideal scenario, the analysis should have been done using the whole volumetry of each viral inclusion, and using live images quantified over time that is yet to be reported.

## ACKNOWLEDGEMENTS

This project has received funding from the European Research Council (ERC) under the European Union’s Horizon 2020 research and innovation programme (grant agreement No. 101001521). Salary support from FCT : T.A.E, D.B., V.M. are funded by PhD fellowships (PD/BD/128436/2017, PD/BD/148391/2019 and UI/BD/152254/2021) and S.V.C by D.L. 57.

## AUTHOR CONTRIBUTIONS

T.A.E., S.V.C, S.S., D.B., I.B., V.M., F.F., M.A, M.S., M.J.A designed and executed experiments and analysed the data. T.A.E AND M.J.A wrote the manuscript. M.J.A. initiated and designed the overall project. S.S., I.B., and M.S. designed, performed, and analysed the experiments of whole proteome solubility assay and T.A.E, D.B., S.V.C, F.F., M.A, M.J.A designed and executed experiments to validate hits. D.B., and M.J.A designed, performed, and analysed animal experiments. T.A.E., V.M., M.J.A designed, performed, and analysed the framework on how the thermodynamic variables influence biophysical parameters and implemented the framework for analyses. T.A.E, S.V.C, M.J.A., designed, performed, and analyzed all live cell analyses. M.S and M.J.A. obtained funding for the study. All authors reviewed the manuscript.

## INCLUSION OF DIVERSITY

One or more of the authors of this paper self-identifies as an underrepresented ethnic minority in science.

## STAR METHODS

Detailed methods are provided in the online version of this paper and include the following:

### KEY RESOURCES TABLE

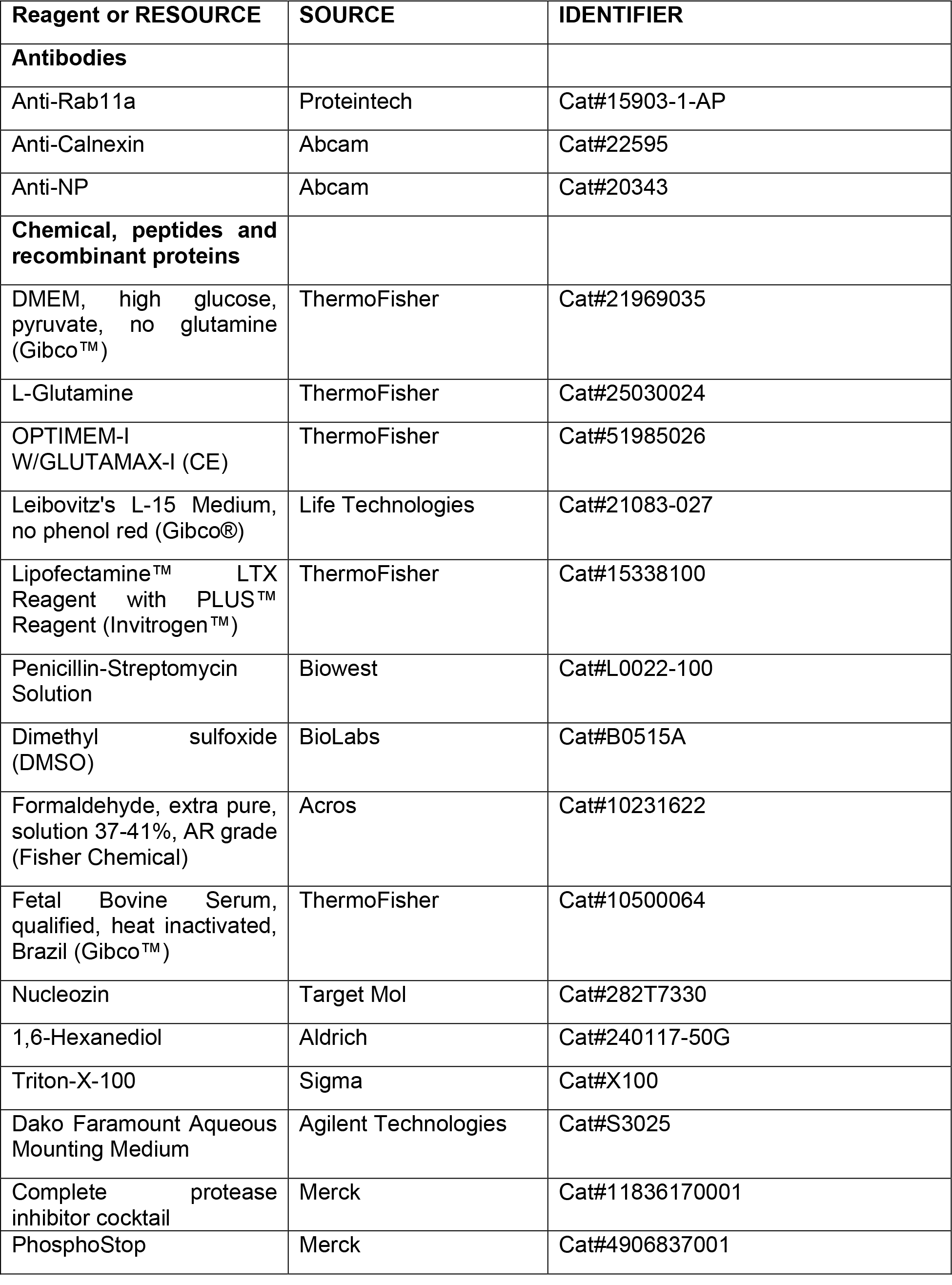
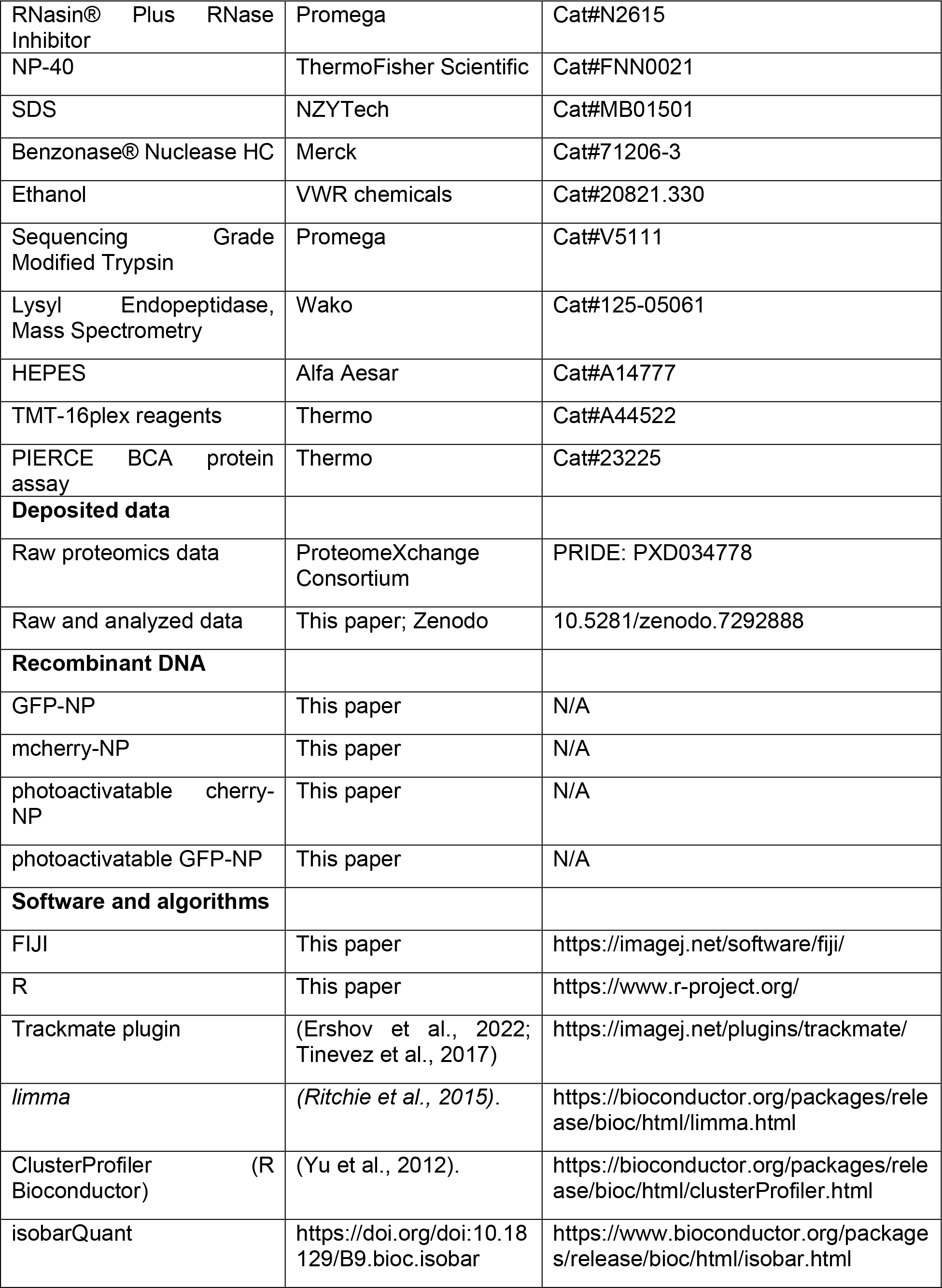
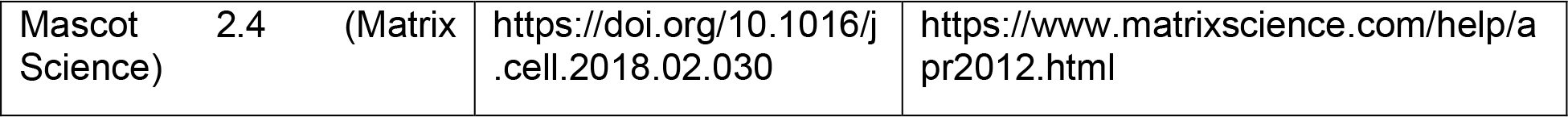

### RESOURCES AVAILABILITY

#### Lead contact

Further information and requests for resources and reagents should be directed to and will be fulfilled by the lead contact, Maria Joao Amorim.

#### Materials availability

This study did not generate new unique reagent.

#### Data availability and code availability

- The mass spectrometry proteomics data have been deposited to the ProteomeXchange Consortium via the PRIDE (PubMed ID: 34723319) partner repository with the dataset identifier PXD034778.

- Reviewer account details:
- Username:
- Password: BprURfLw
- All computer code or algorithm used to generate the results reported in the paper are available at 10.5281/zenodo.7292888.
- All experimental data shown in Figure 1–8 and Figure S1-2 is available from the corresponding author upon request. Sequences of described viruses are accessible from the NCBI virus under accession number GCF_000865725.1. Source data are provided with this paper in 10.5281/zenodo.7292888.

### EXPERIMENTAL MODEL AND SUBJECT DETAILS

#### Cell lines

GFP-Rab11a-WT and GFP-Rab11a-DN cell lines were produced in-house and characterized in (Vale-Costa et al., 2016), while the human alvelolar basal cell (A549) and epithelial cell Madin-

Darby Canine Kidney (MDCK) were generous gifts from Prof Paul Digard, Roslin Institute, UK. Cells were cultured in Dulbeccós Modified Eaglés Medium (DMEM) supplemented with 10% fetal bovine serum (FBS), 2 mM L-glutamine and 1% Pencillin-Streptomycin. GFP-Rab11a cell lines were cultured/maintained in DMEM supplemented with 1.25 µg/mL Puromycin. Cells were maintained in a humidified incubator at 37°C and 5% v/v atmospheric CO2.

#### Viruses

Reverse-genetics engineered A/Puerto Rico/8/34 virus (PR8 WT; H1N1) was used to infect all cell types and titrated by plaque assay in MDCK cells, while X31 virus (a reassortant virus carrying HA and NA segments from A/Hong-Kong/1/1968 (H3N2) in the background of PR8) was used to infect mice. Infection for live imaging were done at 10 MOI, with viral infections for immunofluorescence at an MOI of 3 or 5.

#### Animals and infection

Female C57Bl/6 mice were used.

#### Ethics statement

All experiments involving mice were performed using 8-week-old littermate C57BL6/6J, female mice under specific pathogen-free conditions at the Instituto Gulbenkian de Ciência (IGC) biosafety level 2 animal facility (BSL-2). Animals were group housed in individually ventilated cages with access to food and water *ad libitum*. This research project was ethically reviewed and approved by both the Ethics Committee and the Animal Welfare Body of the IGC (license reference: A003.2021), and by the Portuguese National Entity that regulates the use of laboratory animals (DGAV – Direção Geral de Alimentação e Veterinária (license references: 0421/000/000/2022, Controlling influenza A virus liquid organelles – LOFLU, funded by the European Research Council). All experiments conducted on animals followed the Portuguese (Decreto-Lei n° 113/2013) and European (Directive 2010/63/EU) legislations, concerning housing, husbandry, and animal welfare.

### METHODS DETAILS

#### Mice infection

Female C57Bl/6 mice were infected with 4000pfu of X31 (A/X31; H3N2) virus for 2 days. At 30min, 1h or 2h before the collection of the lungs, each mouse was intranasally treated with PBS (vehicle) or 2.3mg/mL of nucleozin (Ncz). Then, lungs were collected to determine viral titres by plaque assays (using MDCK infected with a set of serial dilutions from the homogenized lung tissue samples) and for histology processing.

#### Plaque assay

For viral titre measurement, A549 cells were seeded for 24 hrs, infected at MOI of 3 in DMEM supplemented with 2 mM L-glutamine and 1% penicillin/streptomycin and devoid of sera for 45 mins at 37 °C and 5% CO2. The supernatants were subjected to a plaque assay in MDCK cells to calculate the virus titres, as described previously (Matrosovich et al., 2006).

#### Drug treatment

Nucleozin was dissolved in dimethyl sulfoxide (DMSO) and used at a final concentration of 2 µM (immunofluorescence staining and virus titres) or 5 µM (live imaging), while 1,6-Hexanediol was dissolved in DMEM and used at 5 % (w/v).

#### Microscopy and image processing

For immunofluorescence, A549 cells were fixed in 4% paraformaldehyde for 10 mins and permeabilized with triton-X-100 (0.2% (v/v)), incubated in primary antibodies for 1 h at RT, washed (3x) in PBS/1% FBS and finally incubated in Hoechst and Alexa Fluor conjugated secondary antibodies for 45mins at RT. Antibodies used were rabbit polyclonal against Rab11a (1:100; Proteintech, 15903-1-AP), calnexin (1:1000, Abcam, 22595), TRIM25 (1: 100, Abcam, ab167154), and NP (1:1000; gift from Prof Paul Digard), mouse polyclonal against NS1 (Neat, in-house from hybridoma made at the IGC antibody facility), mouse monoclonal against NP (1:1000; Abcam, 20343), Tom20 (1:200; Sigma-Aldrich, WH0009804M1) and Drp1 (1:200; Abcam, ab56788). Secondary antibodies were all from the Alexa Fluor range (1:1000; Life Technologies). Following washing in PBS, cells were mounted with Dako Faramount Aqueous Mounting Medium and single optical sections were imaged with a Leica SP5 live or stellaris confocal microscope using the photon counter mode. For z-stacks image, a spinning disk 3i (Marianas) confocal microscope was used in the super-resolution (CSU-W1, SoRa) mode. Samples were imaged on a 63x oil immersion Nikon objective (NA = 1.4). Using the function sum of slices, stacked images were projected to 2D and inclusion and its cytoplasmic milieu were segmented and analyzed using Lab-custom ImageJ macros and R analytics scripts.

#### Determining inclusion topology and thermodynamics

To determine the total concentration of vRNPs (NP as proxy) transported to the cytoplasm in relation to vRNPs produced in the nucleus 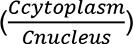, a sum of slices of z-stacked images were used, otherwise, single plane images were analysed for other parameters. We used a custom (Fiji Is Just) ImageJ 2.1.0/1.53p script for image processing using the following pipeline: **(1.)** Segment cell periphery. **(2.)** Segment and remove nucleus from the cell to make the cytoplasm. **(3.)** From the cytoplasm, segment inclusions **(4.)** Analyse the cytoplasm, nucleus, and inclusions for number and topological shape descriptors **(5.)** Using the appropriate segmented region, measure the mean fluorescence intensity (as proxy of concentration) of cell, nucleus, cytoplasm, and cytoplasmic inclusion (*See* **Figure 1***B*).

Using the method published by Riback *et al. 2020* as template, we determined Cdense as the mean fluorescence intensity of the segmented inclusion while Cdilute was extrapolated from remaining cytoplasmic vRNP intensity outside the inclusions. We picked the best approach out of three to measure Cdilute. **(1.)** Use ROIs from randomly selected cytoplasmic areas lacking inclusions. The limitation with this method is that inclusions are highly abundant in the cytoplasm of infected cells and are nearly impossible to manually or automatically draw without selecting regions containing inclusions. **(2.)** Use an enlarged ROI band around the inclusions. This was easy to automate but limited by the overlap with other ROI bands due to the density of IAV inclusions in the infected cell. **(3.)** Use ROI of the entire cytoplasm devoid of viral inclusions. This was easy to automate, lacks overlap with other ROIs and serves as the cleanest strategy when compared to strategy 2 (**Figure S3** A-H). We used strategy 3 to determine the Cdilute.

Partition coefficient (K) and free energy (ΔG) were derived based on Riback *et al.,* 2020 publication; where 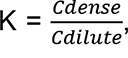, and ΔG = -RT*In*K. Inclusion saturation concentration (Csat) is the threshold Cdilute where inclusion begins to appear (∼ 6hpi) and is calculated as the minimum Cdilute in cells with observable viral inclusions. The change in free energy was normalised to 3hpi (an infection stage with nuclear vRNP staining lacking cytoplasmic inclusions and was represented as ΔΔG = . ΔG – ΔG(3 hpi).

#### Live Imaging, photoactivation

A549 cells were seeded in 8-well glass-bottomed dish (Ibidi) and grown overnight in OptiMEM (37°C, 5% CO2). Cells infected with PR8 at an MOI of 10 were transfected simultaneously with 200ng/µl GFP-NP or cherry-NP plasmid. For photoactivation experiment, a corresponding plasmid of either photoactivatable cherry-NP or photoactivatable GFP-NP was co-transfected with lipofectamine LTX. Cells were imaged using OptiMEM or Leibovitz medium with a 63x oil immersion Nikon objective (NA = 1.4) on Roper TIRF, AiryScan or spinning disk confocal (SoRa) microscopes equipped with temperature (37°C) and CO2 (5%) regulated chamber and stage. Inclusions at a specified region of interest (ROI) was activated by blue light (405 nm laser) at 100% intensity and imaged at 1 frame/ sec for 2 min using 488 nm and 568 nm lasers for GFP and cherry respectively. Photoactivation data were post-processed in FIJI (Image J) using a modified FLAPh algorithm and analysed with a lab-custom R script. Model was obtained using single exponential curve fitting. y = (1-a) + ae^-kt^, a = mobile fraction, K = decay rate constant (per second, s^-1^), t = time (s).

#### Particle tracking and coarsening assay

Trackmate plugin ((Fiji Is Just) ImageJ 2.1.0/1.53p, FIJI) was used to track inclusions at a timescale of 1 s/frame in live imaging samples and XY trajectories were subsequently analysed in a custom R (version 4.1.0) script. Using (FIJI and R), coarsening assay was analysed from time-lapsed tracking of two inclusions, starting from the point they first touch to the point they relax into a rounded puncta with an aspect ratio (AR) of 1.

#### Solubility Proteome Profiling

A549, GFP-Rab11a-WT and GFP-Rab11a-DN cells were mock-infected or infected with PR8 virus between 4 to 16hpi and treated with nucleozin or DMSO. Frozen cell pellets containing 1x10^6^ cells were shipped to Proteomics Core Facility at EMBL, Heidelberg for further sample processing.

Samples for mass spectrometry analysis were prepared as described (Zhang et al., 2022). Briefly, 1x10^6^ cells were resuspended in 100 µl lysis buffer (0.8 % NP-40, 1x cOmplete protease inhibitor cocktail (Roche), 1x PhosphoStop (Roche), 1 U/ml RNAsin (Promega), 1.5 mM MgCl2 in PBS (2.67 mM KCl, 1.5 mM KH2PO4, 137 mM NaCl, and 8.1 mM NaH2PO4, pH 7.4). The sample aliquot for total proteome was incubated directly with benzonase on ice, while the sample aliquot for the soluble proteome was spun down at 100,000 g at 4 °C for 20 min. The supernatant was incubated with benzonase. Both total and soluble aliquots were incubated for 10 min with final 1

% SDS. Protein concentration was determined for the total proteome sample and aliquots equal to 5 µg protein were taken for sample preparation for MS analysis. Both soluble and total lysate of each sample was combined in a multiplexing MS experiment.

#### Mass spectrometry sample preparation

Sample preparation for mass spectrometric measurements were performed as described in (Mateus et al., 2020; Sridharan et al., 2019) .

#### Protein digestion and labelling

Protein digestion was performed using a modified SP3 protocol (Hughes et al., 2014; Hughes et al., 2019). 5 µg of proteins (per condition) were diluted to a final volume of 20 µl with 0.5% SDS and mixed with a bead slurry (Sera-Mag Speed beads, Thermo Fisher Scientific) in ethanol) and incubated on a shaker at room temperature for 15 min. The beads were washed four times with 70% ethanol. Proteins on beads were overnight reduced (1.7mM TECP), alkylated (5mM chloroacetamide) and digested (0.2 µg trypsin, 0.2µg LysC) 100 mM HEPES, pH8. On the next day, peptides were eluted from the beads, dried under vacuum, reconstituted in 10 µl of water and labelled with TMT-16plex reagents for one hour at room temperature. The labelling reaction was quenched with 4 µl of 5% hydroxylamine and the conditions belonging to a single MS experiment were pooled together. The pooled sample was desalted with solid-phase extraction after acidification with 0.1 % formic acid. The samples were loaded on a Waters OASIS HLB µelution plate (30µm), washed twice with 0.05% formic acid and finally eluted in 100 µl of 80% acetonitrile containing 0.05% formic acid. The desalted peptides were dried under vacuum and reconstituted in 20 mM ammonium formate. The samples were fractionated using C18-based reversed-phase chromatography running at high pH. Mobile phases constituted of 20 mM Ammonium formate pH 10 (buffer A) and acetonitrile (buffer B). This system was run at 0.1 ml/min on the following gradient: 0% B for 0 – 2 min, linear increase 0 - 35% B in 2 – 60 min, 35 – 85% B in 60 – 62 min, maintain at 85% B until 68 min, linear decrease to 0% in 68 – 70 min and finally equilibrated the system at 0% B until 85 min. Fractions were collected between 2 – 70 min and every 12^th^ fraction was pooled together and vacuum dried.

#### LC-MS-MS measurement

Samples were re-suspended in 0.05% formic acid, 4% ACN in LC-MS grade water and analyzed on Q Exactive Plus mass spectrometer (Thermo Fisher Scientific) connected to UltiMate 3000 RSLC nano system (Thermo Fisher Scientific) equipped with a trapping cartridge (Precolumn; C18 PepMap 100, 5 μm, 300 μm i.d. × 5 mm, 100 Å) and an analytical column (Waters nanoEase HSS C18 T3, 75 μm × 25 cm, 1.8 μm, 100 Å) for chromatographic separation. Mobile phase constituted of 0.1% formic acid in LC-MS grade water (Buffer A) and 0.1% formic acid in LC-MS grade acetonitrile (Buffer B). The peptides were loaded on the trap column (30 μl/min of 0.05% trifluoroacetic acid in LC-MS grade water for 3 min) and eluted using a gradient from 2 % to 30 % Buffer B over 103 min at 0.3 μl/min (followed by an increase to 40 % B, and a final wash to 80 % B for 2 min before re-equilibration to initial conditions). The outlet of the LC- system was directly fed for MS analysis using a Nanospray-Flex ion source and a Pico-Tip Emitter 360 μm OD × 20 μm ID; 10 μm tip (New Objective). The mass spectrometer was operated in positive ion mode. The spray voltage and capillary temperature was set to 2.2 kV and 275°C respectively. Full-scan MS spectra with a mass range of 375–1,200 m/z were acquired in profile mode using a resolution of 70,000 (maximum fill time of 250 ms or a maximum of 3e6 ions (automatic gain control, AGC)). Fragmentation was triggered for the top 10 peaks with charge 2–4 on the MS scan (data-dependent acquisition) with a 30-s dynamic exclusion window (normalized collision energy was 30), and MS/MS spectra were acquired in profile mode with a resolution of 35,000 (maximum fill time of 120 ms or an AGC target of 2e5 ions).

#### Protein identification and quantification

The MS data was processed as described in (Sridharan et al., 2019). Briefly, the raw MS data was processed with isobarQuant (and identification of peptides and proteins was performed with Mascot 2.4 (Matrix Science) against a database containing *Homo sapiens* Uniprot FASTA ((proteome ID: UP000005640, downloaded on 14 May 2016) and Influenza A virus (strain A/Puerto Rico/8/1934 H1N1, proteome ID: UP000009255) along with known contaminants and the reverse protein sequences (search parameters: trypsin; missed cleavages 3; peptide tolerance 10 ppm; MS/MS tolerance 0.02 Da; fixed modifications included carbamidomethyl on cysteines and TMT16plex on lysine; variable modifications included acetylation of protein N-terminus, methionine oxidation and TMT16plex on peptide N-termini).

#### Mass spectrometry data analysis and normalization

All MS data analysis was performed using R studio (version 1.2.1335 and R version 3.6.1). Data normalization of NP40- and SDS- derived proteomes was performed with *vsn* (Huber et al., 2002). The overall signal sum intensities distributions from all TMT channels of all replicates were corrected for technical variations.

#### Differential analysis of protein abundance

The log2 transformed *vsn* normalized SDS-derived signal sum intensities of proteins from different samples were analysed for differential abundances using *limma (Ritchie et al., 2015)*. Proteins with |log2(fold change) | > 0.5 and adjusted p-value (Benjamini Hochberg) < 0.1 were considered significantly changed.

#### Differential analysis of protein solubility

Solubility is defined as the ratio of NP40- and SDS- derived abundances of proteins. This ratio was computed for all proteins measured in a dataset. The log2 transformed protein solubility was compared between different conditions (time points of infection or different cell line at 12 hours post infection) using *limma*. Proteins with |log2(fold change) | > 0.5 and adjusted p-value (Benjamini Hochberg) < 0.1 were considered significantly changed.

#### Gene ontology over representation analysis

Differential abundant or soluble human proteins from infection time course or different cell line datasets were used for GO term “Biological processes” and/or “Cellular Compartments” overrepresentation analysis using clusterProfiler (R Bioconductor) (Yu et al., 2012). All identified proteins in each dataset served as the background. Standard settings were used for representing enriched GO terms (p-value cutoff: 0.05, Benjamini-Hochberg procedure for multiple testing adjustment and q-value cutoff of 0.2).

### QUANTIFICATION AND STATISTICAL ANALYSIS

After testing for homogeneity of variance, homogenously distributed data were assessed by parametric test using One-way ANOVA, followed by Tukey multiple comparisons of means. In contrast, non-homogenous data were assessed by non-parametric test with statistical levels determined after Kruskal-Wallis Bonferroni treatment. Alphabets above each boxplot represents the statistical differences between groups. Same alphabets indicate lack of significant difference between groups while different alphabets infer a statistically significant difference at α = 0.05.

## SUPPLEMENTAL INFORMATION

This section contains extended material – Movie 1-10, Tables S1-3 and Figure S1-S3, - for the manuscript “Rules for hardening influenza A virus liquid condensates” from the authors Temitope Akhigbe Etibor, Sílvia Vale-Costa, Sindhuja Sridharan, Daniela Brás, Isabelle Becher, Victor Mello, Filipe Ferreira, Marta Bebiano Alenquer, Mikhail Savitski and Maria João Amorim.

## Supplementary Videos

All videos were acquired at the speed of 1second/frame.

**Movie S1:** Fusion and Fission dynamics of IAV inclusions with endogenous Rab11a.

A549 cells expressing endogenous Rab11a were PR8 infected and co-transfected with cherry- NP for 16 hrs and subsequently monitored for inclusion fusion and fission events by live imaging.

**Movie S2:** Fusion and Fission dynamics of IAV inclusions overexpressing Rab11a.

Cell lines overexpressing Rab11a-WT were PR8 infected and co-transfected with cherry-NP for 16 hrs, after which fusion and fusion dynamics were monitored by live imaging.

**Movie S3:** Coarsening assay in liquid inclusions

Liquid-like DMSO-treated inclusions formed in post infected and GFP-NP co-transfected A549 cells were quantified for their ability to coarsen by live imaging. We present 2 videos a-b, showing an entire cell (a) or an inlet with an example of fusion event (b).

**Movie S4:** Coarsening assay in hardened inclusions

Hardened nucleozin-induced inclusions formed in post infected and GFP-NP co-transfected A549 cells were quantified for their inability to coarsen by live imaging. We present 2 videos a-b, showing an entire cell (a) or an inlet with an example of fusion event (b).

**Movie S5:** Fluorescence loss after photoactivation (FLAPh) in liquid inclusions.

PR8 infected A549 was co-transfected with cherry-NP and photoGFP-NP and treated with DMSO for posterior photoactivation of viral inclusions (with blue light, 405 nm) at a region of interest (ROI). Fluorescence loss in the ROI was monitored over time as vRNPs (NP, as a proxy) were transferred from the activated zone to the inactivated region. We present 3 videos a-d, the separate channels for cherry-NP (a), photoGFP-NP (b), the merged video (c).

**Movie S6:** Fluorescence loss after photoactivation (FLAPh) in hardened inclusions.

PR8 infected A549 was co-transfected with photoGFP-NP and cherry-NP and treated with nucleozin for posterior photoactivation of viral inclusions (with blue light, 405 nm) at a region of interest (ROI). Fluorescence loss in the ROI was monitored over time as vRNPs (NP, as a proxy) were transferred from the activated zone to the inactivated region. We present 3 videos a-c, the separate channels for cherry-NP (a), photoGFP-NP (b), the merged video (c).

**Movie S7:** Fluorescence recovery after photobleaching (FRAP) in liquid inclusions

PR8 infected A549 co-transfected with GFP-NP and cherry-NP were treated with nocodazole (Noc) and subsequently photobleached (488 nm) at a region of interest (ROI) in the centre of an inclusion to follow the internal rearrangement by live imaging. We present 2 videos a-b, showing an entire cell (a) or an inlet with an example of fusion event (b).

**Movie S8:** FRAP in hardened inclusions.

PR8 infected A549 co-transfected with GFP-NP and cherry-NP were treated with both nocodazole and nucleozin and subsequently photobleached (488 nm) at a region of interest (ROI) in the centre of an inclusion to see if internal rearrangement occurred during live imaging. We present 2 videos a-b, showing an entire cell (a) or an inlet with an example of fusion event (b).

**Movie S9:** Coarsening assay of Rab11a-DN cells treated with DMSO but lacking inclusions.

GFP-Rab11a-DN cell lines lacking viral inclusion and coarsening events upon infection and transfection (cherry-NP). We present 4 videos a-d, the separate channels for cherry-NP (a), photoGFP-NP (b), the merged video (c).

**Movie S10:** Coarsening assay of hardened viral inclusions formed by treating Rab11a-DN with nucleozin.

GFP-Rab11a-DN cell lines transfected (with cherry-NP) and PR8-infected display a solid-like coarsening behaviour when treated with nucleozin. We present 4 videos a-d, the separate channels for cherry-NP (a), photoGFP-NP (b), the merged video (c).

## Supplementary Tables

Table S1: Topology, thermodynamics and material properties of IAV inclusions. After PR8 infection (1 MOI) inclusions were subjected to thermal changes (Sheet 1) and live imaging for inclusion tracking (Sheet 5), fusion dynamics (Sheet 6) and FLAPh (Sheet 7), or observed for different hours post infection (hpi) as vRNP concentration increases (Sheet 2) or with overexpressed Rab11 (Sheet 3), or nucleozin treatment (Sheet 4). The data summary of inclusion topology and thermodynamics are listed in this table in sheet 1-4. Mice was infected with X31 and treated with PBS or nucleozin for the analysis of number of inclusion and topology in lung slices (Sheet 8).

Table S2: Differential analysis of protein solubility changes before and after nucleozin treatment at 12 hours post infection in WT and Rab11a-DN cell lines PR8-infected Rab11a-WT and Rab11a-DN cells were treated with either DMSO (vehicle) or 5 µM of nucleozin for 1 hour. The protein solubility changes upon nucleozin (or DMSO) treatment in RAB11a-WT and Rab11a-DN cells is listed in this table.

Table S3: Differential analysis of protein abundance changes before and after nucleozin treatment at 12 hours post infection in WT and Rab11a-DN cell lines PR8-infected Rab11a-WT and Rab11a-DN cells were treated with either DMSO (vehicle) or 5 µM of nucleozin for 1 hour. The protein abundance changes upon nucleozin (or DMSO) treatment in RAB11a-WT and Rab11a-DN cells is listed in this table.

## Supplementary Figures

**Figure S1.**
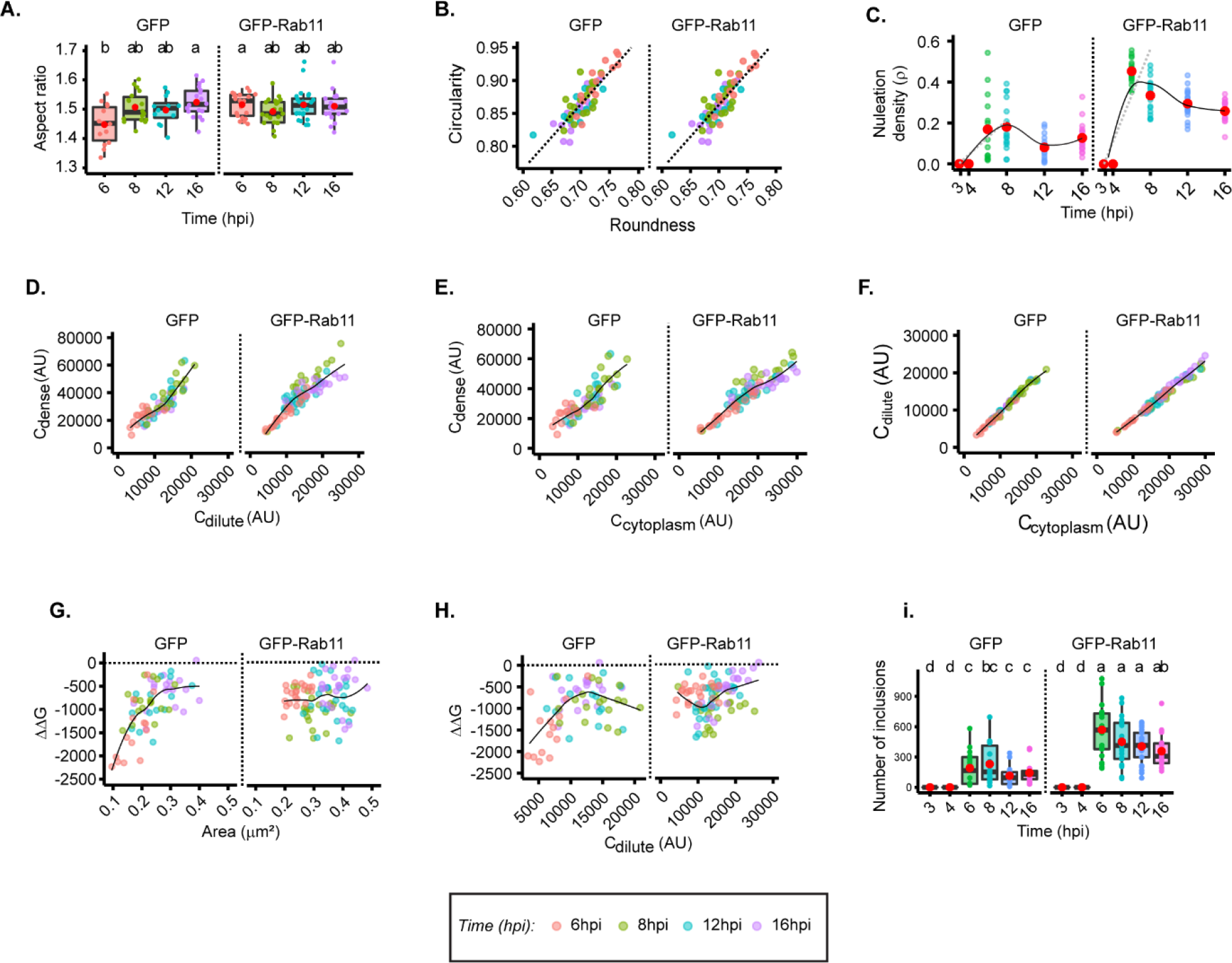
Change in vRNP and Rab11a concentration modestly alter inclusions properties. (A - H) A549 cells stably expressing GFP, or Rab11a-WT, as indicated, were infected at a MOI of 3 with PR8 virus and, at the indicated time points, were fixed, and analysed by immunofluorescence using antibody against NP (as a proxy for vRNPs). The cytoplasmic vRNP concentration increases with time of infection (hpi) and was used as a proxy for cytoplasmic vRNP concentration changes. Each dot is the average value of measured parameter within or outside IAV inclusions per cell, while the continuous black lines are non-linear fitted models for all data. Above each boxplot, same letters indicate no significant difference between them, while different letters indicate a statistical significance at α = 0.05 using one-way ANOVA, followed by Tukey multiple comparisons of means for parametric analysis, or Kruskal-Wallis Bonferroni treatment for non-parametric analysis. Abbreviations: AU, arbitrary unit. (A) Boxplot of inclusion aspect ratio at different hpi. *P* = 0.033422; Kruskal Wallis Bonferroni treatment. (B) Scatter plot of inclusion circularity versus roundness at different time post infection (hpi). (C) Dot plot and model depicting nucleation density (ρ, µm^-2^) over time of infection (hpi). *P* = 0.001; Kruskal Wallis Bonferroni treatment. (D) Scatter plot of Cdense (AU) versus Cdilute (AU) at different hpi. (E) Scatter plot of Cdense (AU) and Ccytoplasm (AU). (F) Scatter plot of Cdilute (AU) versus Ccytoplasm (AU) with time of infection. Coloured lines are non-linear fitted models of the data points in the graph (G - H) Conditions were normalized to an infection state without IAV inclusions (3 hpi) that is indicated by the dashed black line. (G) Scatter plot of ΔΔG (J.mol^-1^) relative to 3 hpi versus area of inclusion. (H) Scatter plot of ΔΔG versus Cdilute (AU) with time of IAV infection. (I) Boxplot of inclusion number per cell at different hpi. *P* = 0.001; Kruskal Wallis Bonferroni treatment.

**Figure S2.**
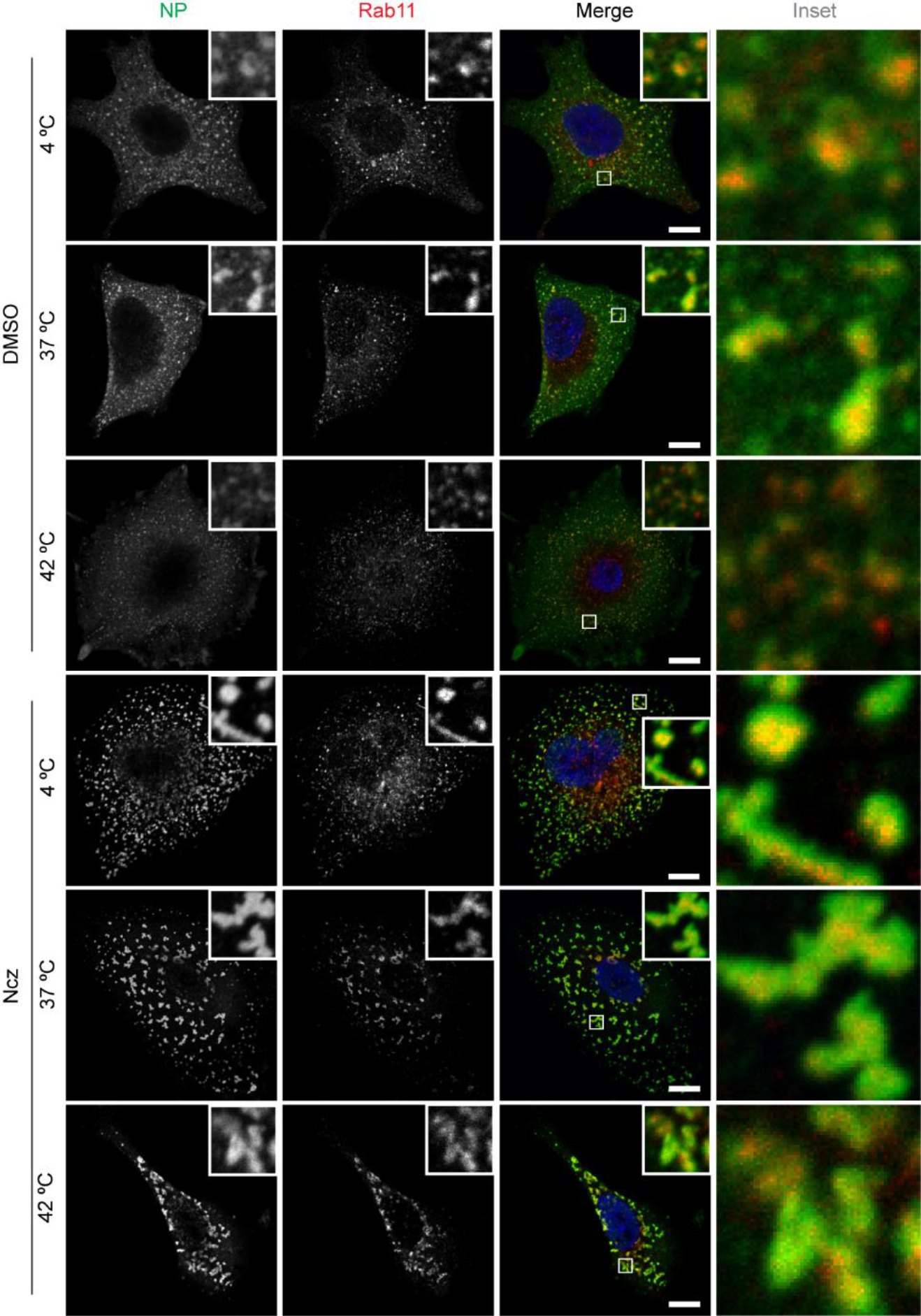
Hardened inclusions are thermally stable. A459 cells were infected with PR8 at a MOI of 3. At 7.5 hpi the infected cells were treated with 5 µM Ncz or DMSO for 30 mins at 37°C before being subjected to thermal stress at 4°C, 37°C and 42°C for 20mins and fixed for immunofluorescence analysis by staining with antibody against NP (green), Rab11 (red) and nucleus (blue). Representative images with Scale Bar = 10 µm.

**Figure S3.**
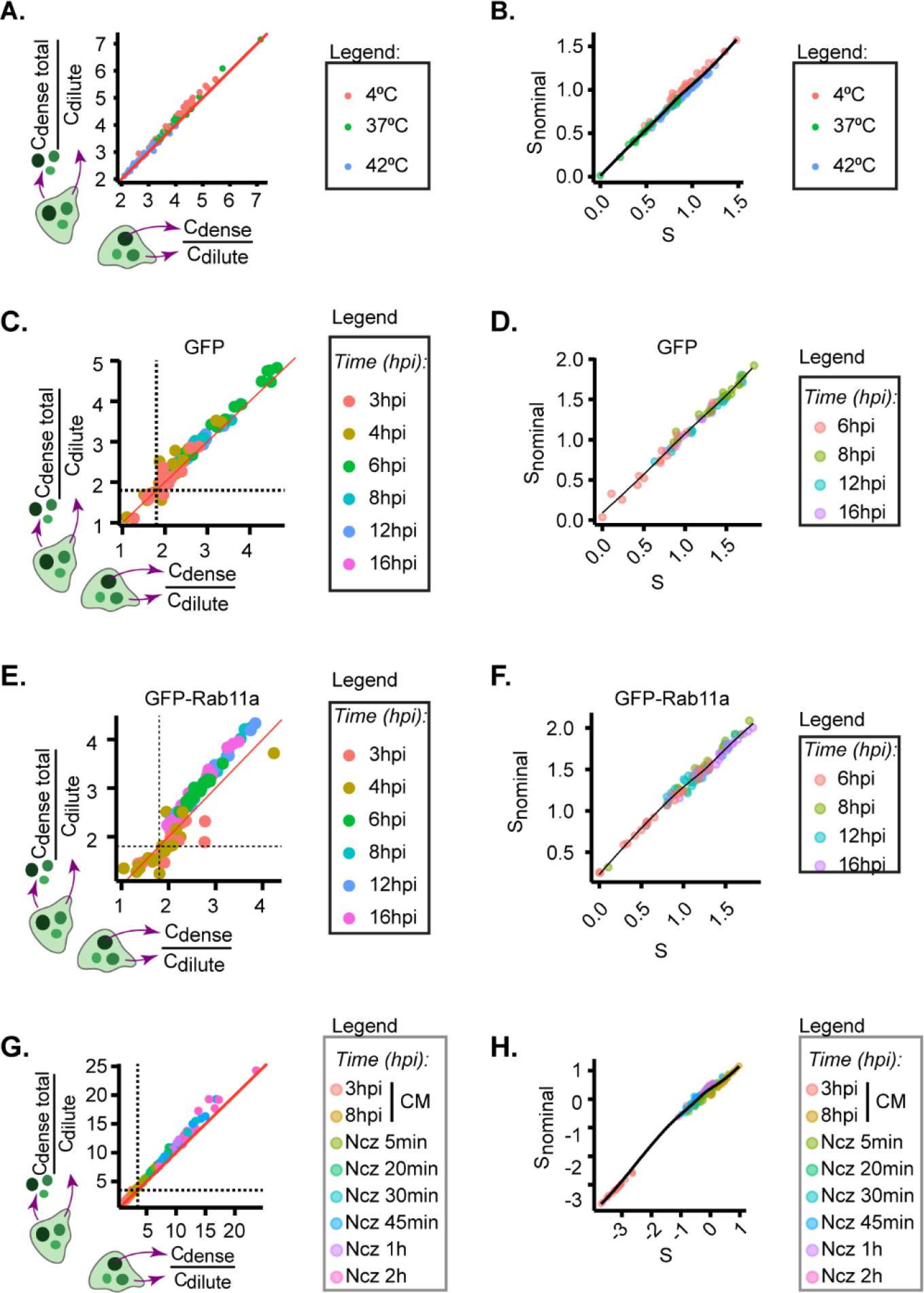
Validation of method analysing thermodynamics parameters. A549 cells expressing (A – D, G – H) endogenous levels of Rab11a or (E - F) over expressing Rab11a were infected at a MOI of 3 with PR8 virus for (A – B, G - H) 8 h before incubating the cells at the indicated (A - B) temperatures, (G - H) Ncz residence time or (C - F) at the indicated timepoints. After this, the cells were fixed, and analysed by immunofluorescence using antibody against NP (as a proxy for vRNPs). Each dot is the average value of measured parameter within or outside IAV inclusions per cell, while the continuous black lines are non-linear fitted models for all data. (A, C, E, G) are the scatterplots comparing image segmentation strategies to calculate partition coefficient and extrapolate the free energy (*see Methods*) while (B,D,F,H) is a scatter plot comparing methods for calculating the degree of supersaturation.

